# The reduction of nuptial gifts in sclerosomatid Opiliones coincides with increased behavioural sexual conflict

**DOI:** 10.1101/2024.06.13.598848

**Authors:** Tyler A. Brown, Emily Marinko, Mercedes Burns

## Abstract

Nuptial gifts serve to increase donor fitness through a variety of pathways. Some nuptial gifts deliver benefits similar to those of antagonistic male behaviours: functioning to secure additional copulations, increase sperm transfer or storage, or increase paternity share. These commonalities may result in evolutionary transitions between solicitous and coercive strategies, wherein behavioural sexual conflict could function to secure mates in lieu of nuptial gifts. In temperate leiobunine harvesters (Arachnida: Opiliones), nuptial gifts have been repeatedly lost, resulting in two primary mating syndromes: an ancestral, sacculate state in which males endogenously produce high-investment nuptial gifts and females lack pregenital barriers, and a derived, nonsacculate state in which females have pregenital barriers and males produce significantly reduced, low-investment nuptial gifts. In this study, we investigated whether behavioural sexual conflict compensates for reduced nuptial gifts in nonsacculate harvesters by comparing the intensity of pre-, peri-, and postcopulatory mating behaviour between the nonsacculate species *Leiobunum vittatum* and *L. euserratipalpe* and the sacculate species *L. aldrichi* and *L. bracchiolum*. We additionally sought to establish an automated behavioural analysis pipeline by developing analogues for metrics traditionally scored manually. Our results revealed significantly different, potentially coercive, behaviour in nonsacculate species, indicating that the loss and reduction of pre- and pericopulatory nuptial gifts may contribute to increased behavioural antagonism. Mating behaviour also differed significantly between *L. vittatum* and *L. euserratipalpe*, indicating there are multiple suites of potentially antagonistic behaviours. Together, these results suggest that multiple behavioural strategies may be effective substitutes for nuptial gifts in leiobunine Opiliones, although the mechanisms through which male fitness is increased requires further research.

**Highlights:**

- Temperate Opiliones have repeatedly evolved reduced nuptial gift investment
- Reduced-gift (nonsacculate) species possess morphology indicative of antagonism
- We tested if mating behaviour differed between nonsacculate and sacculate species
- Nonsacculates displayed significantly higher behavioural antagonism than sacculates
- Nonsacculate species differed in their suites of antagonistic behaviours

## Introduction

### Nuptial gift background

Animals employ a multitude of tactics to increase their fitness during mating interactions, including direct intrasexual competition, mate choice, elaborate signals, and physical or chemical coercion (Buchinger & Li 2023; Davranoglue et al. 2023; Edward & Chapman 2011; Órfão et al. 2023; Rosvall 2011; Schaefer & Ruxton 2015; Shuker & Kvarnemo 2021). Of these myriad strategies, nuptial gifts are among the most unique in their almost altruistic appearance. Nuptial gifts are any materials, beyond gametes, which are provided by a donor to a recipient before or during copulation in order to increase the donor’s fitness (Gwynne 2008; Lewis & South 2012; Lewis et al. 2014). These gifts might be consumed orally or received internally during copulation and can range from exogenous materials (such as captured prey wrapped in silk) to endogenous materials (such as proteinaceous fluids) (Gao et al. 2019; Lewis & South 2012). The fitness consequences to the recipient remains intentionally unspecified under this definition, and extensive research shows nuptial gifts can range from being highly beneficial (Albo et al. 2017; Brown & Barry 2016; Gwynne 2008; Lehmann & Lehmann 2016; O’Hara & Brown 2021; Toft & Albo 2015; Voigt et al. 2005; Welke & Schneider 2011) to neutral (Martínez Villar et al. 2021) to highly costly (Arnqvist & Rowe 2005; Sakaluk et al. 2019; Sirot et al. 2015; Wolfner 2002). Predicting these consequences, however, can be difficult due to the highly variant nature of many nuptial gifts, which often vary with source, delivery mechanism, and both within and between individuals (Kahn et al. 2018; Lewis & South 2012; Wedell 1994). Gifts may also fluctuate with environmental conditions, rearing conditions, donor nutrition, or competition intensity (Fox et al. 2006; Ghislandi et al. 2018; Jarrige et al. 2015; Křemenová et al. 2021; McPherson et al. 2022; Rapkin et al. 2016), and males may even adjust gifts based on recipient traits (Jarrige et al. 2015). Donor fitness benefits have long been debated with little conclusive support for a single hypothesis (Arnqvist & Rowe 2005; Gwynne 2008; Lewis & South 2012; Lewis et al. 2014; Sakaluk 2000; Thornhill 1976; Vahed 1998, 2007). Lewis and South (2012) present a revised approach to describing nuptial gift benefits, suggesting eight non-exclusive benefits through which donor fitness may be increased (increasing mate attraction, copulation success, insemination success, sperm transfer, sperm storage, paternity share, female fecundity, or offspring fitness). Behavioural modifications can serve many of these functions, often increasing copulation and insemination success, sperm storage and transfer, and paternity share (Lewis & South 2012; Gwynne 2008; Arnqvist & Rowe 2005). .

The presence of beneficial nuptial gifts does not necessarily signal an interaction lacking conflict (Arnqvist & Rowe 2005; Gwynne 2008; Lewis & South 2012; Sirot et al. 2015). Mating interactions can be costly to females and males – direct physical or physiological harm, resistance costs, investment in gametes or nuptial gifts, or simple energy expenditure and predation risks can each render mating a potentially costly encounter (Arnqvist & Rowe 2005; Boulton et al. 2018; del Castillo & Gwynne 2007; Kokko & Jennions 2014; Lehtonen et al. 2012; Małek et al. 2019; Perry & Tse 2013; Prokop & Okrouhlik 2021; Sentenská et al. 2020; Tatarnic et al. 2014). The costs of producing or procuring nuptial gifts often lead to selective pressures to maximise gift efficiency and efficacy through various strategies, including deceiving females or plastically altering gifts in response to female quality (Albo et al. 2019; Beyer et al. 2021; Ghislandi et al. 2018; Martínez-Villar et al. 2023; Pandulli-Alonso et al. 2021; Solano-Brenes et al. 2021; Stålhandske 2002).

Trade-offs between reproductive tactics, such as those between nuptial gifts and extended clasping or deception, have been increasingly documented across arthropod species (del Castillo & Gwynne 2007; Lehmann et al. 2016; Liu et al. 2015; Miller et al. 2019; Simmons & Emlen 2006; Simmons et al. 2017; South et al. 2011; Yamane et al. 2010). Vahed et al. (2014) demonstrate this, providing convincing evidence that prolonged clasping time can function as a substitute for nuptial gifts. This has subsequently been shown in other orthopterans, suggesting that these nuptial gifts function in some capacity to increase sperm transfer (Gwynne et al. 2008; Lehmann et al. 2016; Vahed et al. 2014). This is also evident in *Neopanorpa* scorpionflies, in which the sole species that has lost nuptial gifts (*N. longiprocessa*) has evolved enhanced clasping structures (Tong & Hua 2019; Zhong & Hua 2013). In this species, mating interactions are characterised by intense female struggle while males attempt to clasp prior to intromission. *Leiobunum* Opiliones show a highly similar pattern of ancestral nuptial gifts and derived nuptial gift reduction in conjunction with enhanced clasping traits, although to date, behavioural studies that directly compare multiple species are lacking, and to our knowledge this is the first study to do so.

### Study Species

North American sclerosomatid harvesters (Opiliones: Sclerosomatidae), also known as “daddy-longlegs”, form a widespread family of arachnids with a mating system ideal for studying the role of nuptial gifts in reproductive behaviour (reproductive biology reviewed in Machado & Burns 2023). *Leiobunum* species possess highly variable genitalia in both females and males, with genital morphology being one of the primary methods of species identification in both sexes (Shultz 2018). All *Leiobunum* females have a moveable, “hinged” genital operculum that covers their genital chamber and must be opened prior to intromission (Shultz & Pinto-da-Rocha 2007; Shultz 2018). Some species have additional pregenital barriers and sclerotization on the operculum which engages an interior sternum, effectively locking the genital opening, but these only occur in species which have also lost nuptial gift sacs (Burns et al. 2013; Shultz 2018). Moveable female barriers are often indicative of sexual conflict, and Eberhard (2010) notes that these barriers may be particularly likely to occur in systems undergoing sexually antagonistic coevolution (though they are not entirely impossible under cryptic female choice). Species in which males have lost nuptial gift sacs also have unique physical and mechanical penis properties, with some species possessing robust penises capable of applying significant force to female structures and other species occupying an intermediary state (Burns & Shultz 2015, 2016). Burns and colleagues have described this repeated correlated evolution of species-specific female and male reproductive morphology, revealing two primary reproductive syndromes (Burns et al. 2012, 2013): 1) An ancestral, sacculate state in which males have terminal nuptial gift sacs on their penises and females do not have pregenital barriers; and 2) a derived, nonsacculate state in which males do not have terminal nuptial gift sacs on their penises and females have pregenital barriers.

Both sacculate and nonsacculate *Leiobunum* harvesters produce nuptial gifts to some degree, although the presence of terminal nuptial gift sacs increases the quantity transferred to females during mating interactions and allows for precopulatory transfer (Machado & Burns 2023; Machado & Macías-Ordóñez 2007). Sacculate females receive nuptial gift fluid before and during copulation, potentially allowing for better precopulatory assessment of males (as expected under cryptic female choice; Eberhard 1996), while nonsacculate females can only receive it after committing themselves to copulation (Figure 1). In addition to these morphological differences, sacculate and nonsacculate species also differ in the biochemistry of their nuptial gifts; sacculate species have nuptial gifts with significantly higher proportions of essential amino acids compared to nonsacculate gifts, and gift composition corresponded to the presence/absence of gift sacs rather than phylogeny (Kahn et al. 2018). Essential amino acids are those that cannot be produced by an organism and must come from their food, likely making them more costly for a male to include in a gift (Kahn et al. 2015). These are also likely particularly valuable for females to receive due to their contribution towards oogenesis (Jarrige et al. 2015; Kahn et al. 2018; Wheeler 1996). This transition from high- to low-value nuptial gifts correlates with morphological transitions between sacculate and nonsacculate states and is consistent with the expectations of a system high in sexual conflict, in which males reduce investment in a highly-costly gift in favour of coercion. Alternatively, it is possible that the reduction of nuptial gifts is instead related to a shift in female preferences, with nonsacculate females evaluating males on behavioural cues rather than biochemical or nutritional cues (Eberhard 1996).

**Figure 1:**
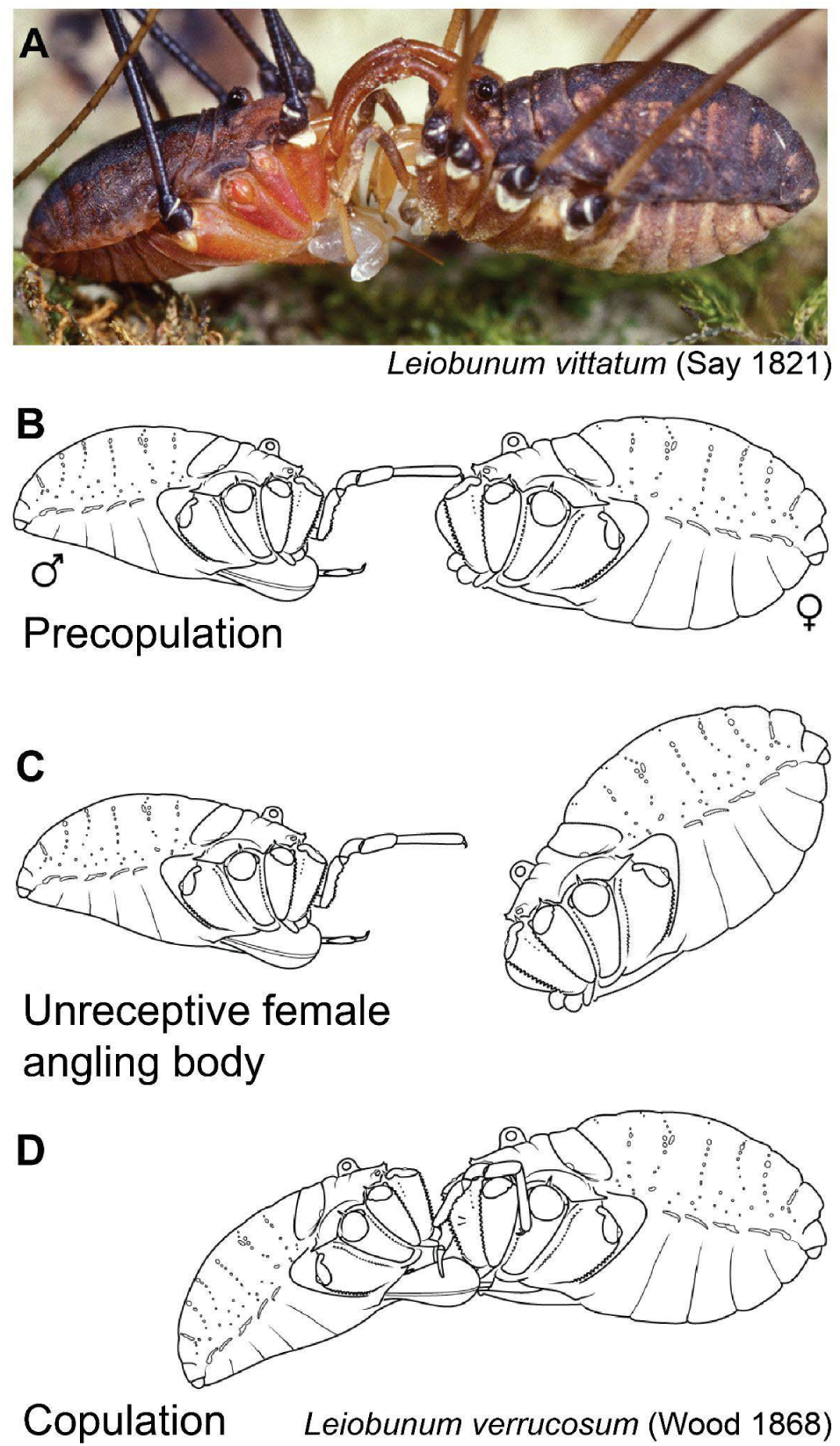
Primary mating phases and body orientations of *Leiobunum spp.* **A)** Male (left) *L. vittatum* clasping female (right) precopulation. Photo courtesy of Joe Warfel. **B)** Precopulatory phase, during which consumption of precopulatory nuptial gift (when present in the species) occurs. **C)** Precopulatory posture of an unreceptive female. Female’s anterior is angled downwards to prevent male access to the ventrally located genital operculum. **D)** Copulatory phase, during which consumption of pericopulatory nuptial gift occurs. **B-D)** Images are semi- diagrammatic and the legs are not pictured to increase clarity (*L. verrucosum*, adapted from Figure 1 in Burns et al. 2013).

The *L. vittatum* and *L. calcar* species-groups have species with pedipalps that are structurally enhanced to facilitate more effective clasping (Burn et al. 2013; Ingianni et al. 2011).

Notably, these exaggerated features only occur in nonsacculate species with female pregenital barriers, though they do not contact these structures during mating. Enhanced-clasping pedipalps evolved independently in these groups, and in all instances this occurred after the transition to a nonsacculate state (Burns et al. 2013). These clasping-structure modifications may allow males to clasp and mate with females for longer than is ideal for the female (and therefore circumventing potential female preferences), and longer intromission periods are highly associated with increased sperm transfer, mate guarding, and sexual conflict in a number of species (Arnqvist & Rowe 2005). Alternatively, the reduction of nuptial gifts in nonsacculate species may be compensated for with an extended period of evaluation and interaction that is facilitated by better clasping structures, and clasping alone is not necessarily indicative of conflict. Critical information may be conveyed during this process, allowing an organism to evaluate a potential mate. Genital clasping by males may provide stimulation to females (Eberhard 2010), and some species with exaggerated genital clasping structures do not necessarily have high levels of sexual conflict (Briceño & Eberhard 2009; Briceño et al. 2015; Myers et al. 2016; Peretti & Aisenberg 2015). However, non-genital clasping structures, such as the pedipalps of Opiliones, are potentially more indicative of sexual conflict (Arnqvist & Rowe 2005; Rowe & Arnqvist 2012).

Although there is morphological and chemical evidence indicative of a potential shift towards increased conflict, comparative behavioural evidence in the genus is limited. Fowler-Finn and colleagues (2014, 2018, 2019) have thoroughly documented the mating behaviour of the nonsacculate *L. vittatum* as well as several other species, but no direct, comparative, behavioural work between sacculate and nonsacculate species has been conducted. The presence of two nuptial gifts in some *Leiobunum* species is unique, and the multiple independent transitions to a derived, nonsacculate state provide an ideal system for studying the role of behaviour in the reduction of nuptial gifts and whether sexual conflict may compensate for this change. In this study, we tested whether nonsacculate leiobunine species displayed significantly different reproductive behaviour than sacculate *Leiobunum* species. We hypothesised that the reduction of nuptial gifts in nonsacculate species of *Leiobunum* would correspond with significantly different mating behaviour when compared to sacculate species. Further, we expected nonsacculate males to compensate for this reduction in gift value with increased coercive behaviour and overall higher levels of sexual conflict. Alternatively, behavioural changes may be driven by shifts in female preferences and mate assessment pathways. To test this, we performed mating trials using two sacculate and two nonsacculate *Leiobunum* species and compared the intensity and frequency of ostensibly antagonistic behaviour to search for differences between the two reproductive syndromes.

## Methods

### Study Species

Leiobunine harvesters have a broad, Holarctic distribution and require extensive taxonomic revision (Hedin et al. 2012), however, the “late-season” (defined as overwintering as eggs) leiobunines of eastern North America form a well-supported monophyletic group (Burns et al. 2012). We therefore restricted our study species to this clade, within which there are three major phylogenetically-supported groups of harvesters. From these we selected one representative species per group, as well as one species which was not a member of any major group (Table 1; see Figure A1 for the full phylogeny).

**Table 1:** Morphological and biochemical phenotypes of the four species in this study, as well as overall behaviour expected in trials (Burns et al. 2012; Kahn et al. 2018; Shultz 2018). High conflict would be indicated by reduced latency to make a clasping attempt, elevated contact duration, elevated time within one leg span, elevated clasping duration, elevated guarding duration, elevated female body angle, reduced association distance, and reduced pursuit angle. Low conflict would be indicated by the inverse of these metrics.

| Species | Species-group | Penis Morphology | Pedipalp Morphology | Nuptial Gift Chemistry | Behavioral Predictions |
| --- | --- | --- | --- | --- | --- |
| *L. euserratipalpe* | *calcar* | Nonsacculate | Unmodified pedipalps | Low investment | High conflict |
| *L. vittatum* | *vittatum* | Nonsacculate | Male pedipalps specialized for clasping | Low investment | High conflict |
| *L. brachiolum* | *politum* | Sacculate | Unmodified pedipalps | High investment | Low conflict |
| *L. aldrichi* | - | Sacculate | Unmodified pedipalps | High investment | Low Conflict |

In sacculate species, males first use their pedipalps to clasp females face-to-face by the coxae of leg II (Figure 1A and D). The male then extends its penis to the female’s mouth, allowing the female to consume the endogenously-produced precopulatory nuptial gift that accumulates in the terminal sacs after being produced by accessory glands (Figure 1B). The male redirects its penis to the female’s genital operculum, which the female then opens to allow intromission. At this stage, the female consumes the endogenously-produced, pericopulatory nuptial gift directly from the male’s papilla as its mouth aligns with male’s haematodocha (Figure 1D; Burns et al. 2012, 2013, Machado & Macías-Ordόñez 2007). In nonsacculate species, this behavioural sequence proceeds much the same, save for two notable differences: 1) the precopulatory nuptial gift does not occur, and 2) the male may use its penis to pry the female genital operculum open should the female resist copulation (Burns et al. 2012, 2013).

### Study Populations

We collected harvesters as juveniles in the spring and summer of 2021, 2022, and 2023. The exact timing and location of collection depended upon the species, as each species matures at different times, and some are restricted in their geographic range across central Maryland. *Leiobunum vittatum* were collected in June and July of 2021 at Rockburn Branch Park (39.219471, −76.761198) and the University of Maryland, Baltimore County (39.247397, - 76.710568); *L. aldrichi* were collected in June and July of 2021 at Sugarloaf Mountain (39.261630, −77.395560) and Plummer’s Island (39.261630, −77.395560); and *L. euserratipalpe* were collected in April and May of 2022 at Schooley Mill Park (39.166325, −76.961830); and *L. bracchiolum* were initially collected in May and June of 2021 at Rockburn Branch park. Due to a low rate of successful clasps, additional *L. bracchiolum* were collected in May and June of 2023 at the same location to ensure sufficient data for analyses which required successful clasps.

Following collection, we separated harvesters by species and sex and housed them communally at 24° C, 80% humidity, and 12:12 h light:dark periods in 37 cm x 22 cm x 24.5 cm plastic terraria. Each terrarium contained ∼ 2 cm of Eco Earth coconut fibre substrate (Zoo Med), petri dishes with TetraMin Select tropical flakes fish food (Tetra), moist cotton balls, and leaves, sticks, and bark gathered from the collection locations for shelter. All terraria were moistened daily and checked for mould, and all were cleaned weekly. Upon maturation, we maintained harvesters in similar conditions until mating trials were complete, at which point we housed them communally by sex until death.

### Behaviour

Behavioural trials consisted of pairing male and female conspecifics and recording mating behaviour. We conducted all behavioural trials in a 29.21 cm x 29.21 cm acrylic arena lined with white cardstock to reduce reflections and allow for easy cleaning. The arena was housed in a room kept at 24° C and illuminated with white light (leiobunine Opiliones have poor vision and lighting is unlikely to impact mating behaviour (Machado & Macías-Ordόñez 2007; Shultz & Pinto-da-Rocha 2007) and we video recorded interactions dorsally and laterally with Microsoft LifeCam Studio cameras. Following the methods used in studies by Fowler-Finn and colleagues (2014, 2018, 2019), we conducted trials as follows: We first sterilised the arena with 95% ethanol before placing a male and female harvester in the arena. Initially, harvesters were isolated for 5 minutes under transparent plastic containers to allow acclimation to the arena without contact with one another. Following acclimation, we removed the containers and the trial began. Trials lasted for either 20-minutes or until the mating interaction ended. Interactions were considered concluded when contact was broken between individuals and the male made no attempt to chase the female. If the interaction had not ended by the 60-minute mark, the trial was concluded.

We aimed to complete 20 trials per species, though the exact number per species varied due to unexpected deaths and exclusions due to inactivity (Appendix Table A1). Trials were included in analyses if the male made an attempt to hook its pedipalps around the female’s coxae within the first 20 minutes. If no mating attempt was made at this point, we removed the harvesters from the arena, excluded them from future trials, and did not include the trial in analyses. The majority of males (86.36%) made an attempt to clasp within the first 20 minutes, meaning only 12 of 88 potential trials were excluded due to inactivity. The majority of interactions (86.67%) had concluded before the imposed 60-minute limit, with only 11 of 75 trials artificially ended. Interestingly, seven of these 11 trials were with *L. euserratipalpe* pairs.

### Video Analyses

We used the automated tracking program TRex (Walter & Couzin 2021) to track individual identities and coordinates to calculate the distance between individuals, total time individuals were within one male leg span (measured using the longest leg length specific to the male in each trial), and total clasping duration (Table 2). We additionally explored a method to assess the degree to which males followed females between first contact and first mating attempt. We used the centroid coordinates to first calculate male and female trajectory vectors and magnitudes before using these to derive the angle between vectors. This metric describes the “pursuit angle” between male and female movement; values closer to 0 degrees indicate the individuals were moving in the exact same direction, which we would expect if the male were chasing the female along parallel vectors (Table 2). Conversely, values closer to 180 degrees indicate the individuals were moving in the exact opposite direction. Some behaviours, particularly those occurring on the ventral side of the harvesters or those involving legs, could not be reliably scored with TRex and we instead used manual scoring to record contact time, female body angle, instances of investigatory leg tapping bouts and instances of females fleeing males (Table 2).

**Table 2:**
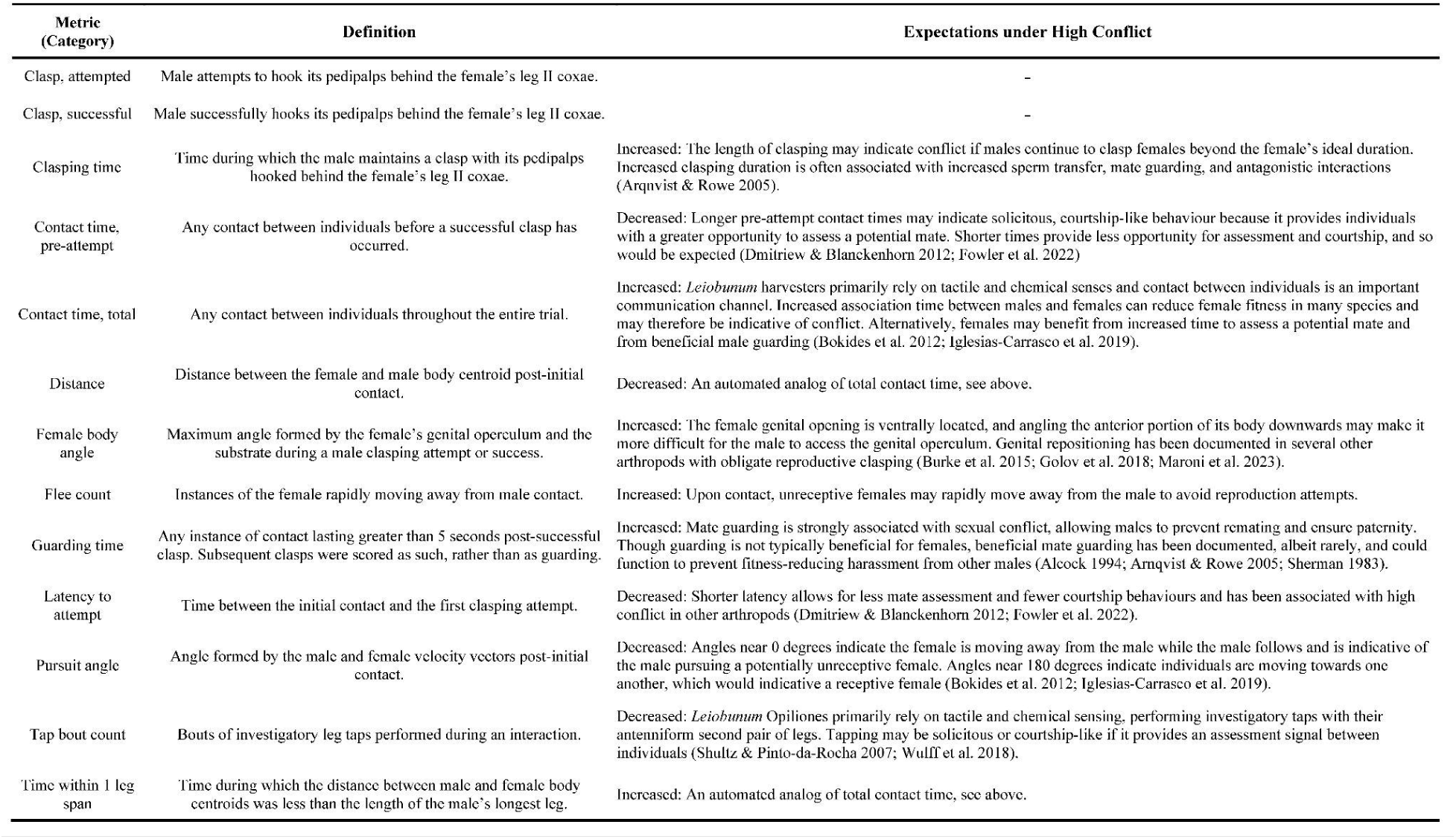
Behavioural metrics recorded during all trials. Rationale for each metric is noted in text with supporting citations.

To ensure TRex performed accurately and matched manual tracking expectations, we additionally scored all trials with JWatcher (Blumstein & Daniel 2007). For all attempted-clasp trials, we scored the total time the harvesters were in contact, the latency between first contact and first mating attempt, the number of female fleeing instances, and the number of “tapping bouts” by the male (Table 2). For successful-clasp trials, we additionally scored the time between first mating attempt and first successful clasp, the total time the individuals were in contact, the total time the male clasped the female as a proxy for intromission time (as per Fowler-Finn et al. 2014, 2018, 2019), and the total time the male guarded the female post-copulation (Table 2). In leiobunine harvesters, mate guarding consists of the male keeping in constant contact with at least one leg grasping the female’s leg; because they primarily rely on tactile sensing using antenniform legs and their vision is particularly poorly-developed, this contact is likely required (Machado & Macías-Ordόñez 2007; Shultz & Pinto-da-Rocha 2007; Fowler-Finn et al. 2018, 2019). We considered post-copulatory contact lasting greater than 5 seconds to constitute mate guarding (Fowler-Finn et al. 2014, 2018, 2019).

Finally, in all attempted-clasp trials, we used the image analysis program FIJI-ImageJ (Schindelin et al. 2012) to measure the maximum angle the female’s genital operculum formed with the substrate of the arena (Figure 1C). Due to the degree to which some mating pairs moved during their interactions, we could not effectively automate this metric to obtain a continuous measure of the angle, thus we elected to use the maximum angle reached during a clasp attempt (Table 2).

### Statistical Analyses

Due to our selected study species being closely related, we first tested for phylogenetic signal across all our behavioural metrics. Previous studies have not found evidence for signal among morphological traits (Burns & Shultz 2015); and significant evidence suggests that behavioural traits are among the least likely to display patterns of phylogenetic signal because of their lability (Blomberg et al. 2003). We imported our behavioural data to R (v4.3.1; R Core Team 2023) and calculated Pagel’s lambda (Pagel 1999) using the *phytools* package (Revell 2012). We used a pruned version (consisting of our four study species) of a maximum clade credibility tree constructed from mitochondrial and nuclear sequences by Burns et al 2013. We found extremely low levels of phylogenetic signal (Likelihood ratio test versus Brownian motion: All *P* > 0.6; Appendix Table A2) and after consideration with previous evidence (Burns & Shultz 2016, 2015; Blomberg et al. 2003) proceeded with frequentist statistical analyses.

Leiobunine harvesters primarily rely on tactile senses; sweeping and tapping their antenniform second pair of legs to investigate their surroundings (Shultz & Pinto-da-Rocha 2007; Willemart & Hebets 2012). We therefore evaluated normality using the Shapiro-Wilk test and followed this with Kruskal-Wallis tests for nonparametric data to determine whether the individuals used in our study differed significantly between species in the number of sensory legs and total legs. Because Opiliones can autotomize their legs, severing them to escape potential predators (Gnaspini & Hara 2007), not all individuals in the study had the same number of total legs or sensory legs. We incorporated this variability by constructing multiple linear regression models with behavioural metrics as response variables (Table 2) and leg count metrics (number of male legs, number of male sensory legs, number of female legs, and number of female sensory legs) as predictor variables. The results of these multiple linear regression models allowed us to determine whether the loss of legs impacted behavioural sexual conflict.

For all behavioural metrics, we used the Shapiro-Wilk test for normality and found at least one species in all but one metric to be non-normally distributed. Due to this, we used the Kruskal-Wallis test to compare the four species for all metrics except the average distance between individuals. For both attempted- and successful-clasp trials this metric was normally distributed after a logarithmic transformation, and so these data were analysed with a one-way ANOVA after transformation. We followed any significant groupwise comparisons with planned pairwise comparisons using Dunn’s multiple comparisons test or Šídák’s multiple comparisons test, where appropriate, between sacculate and nonsacculate species (*L. bracchiolum x L. vittatum*; *L. bracchiolum x L. euserratipalpe*; *L. aldrichi x L. vittatum*; and *L. aldrichi x L. euserratipalpe*). Following inspection of several metrics (pursuit angle, attempt latency, clasping duration, and maximum female body angle), we decided to compare the two nonsacculate species (*L. vittatum* and *L. euserratipalpe*) to one another with post-hoc Dunn’s tests, corrected for multiple comparisons.

We concluded our statistical analyses with multiple logistic regression models to detect which behavioural metrics best predict clasping success. We built models with success/failure to secure a clasp as the response variable and the behavioural metrics as the predictor variables. We constructed a total-species model using all data, a sacculate-species model using only *L. bracchiolum* and *L. aldrichi* data, and species models using species-level data. We could not build nonsacculate-species or species-level models for *L. vittatum* and *L. euserratipalpe* due to each species having only one failed clasp attempt, thus leading to perfect separation in our models.

### Ethical Note

We collected *L. vittatum*, *L. euserratipalpe*, *L. aldrichi*, and *L. bracchiolum* from populations throughout their natural range in central Maryland. The Washington Biologists’ Field Club provided collection permission for Plummer’s Island, while collection permits were not required for any other location. All source populations used for collections were well-established, with the species in question being clearly abundant and unlikely to be affected by our collections. The harvesters were then held in the lab following ASAB/ABS guidelines for the treatment of animals used in behavioural research. Harvesters were reared collectively whenever possible in accordance with their gregarious behaviour in the field. Behavioural trials were generally not harmful to individuals – females experienced harassment from males during trials, but it was likely far less frequent than harassment experienced in natural conditions. Following trials, individuals were maintained in the laboratory until natural death, typically within one to two months.

## Results

### Overall Contact and Association Metrics

We found one significant difference when comparing the number of legs between species (total legs, female *L. bracchiolum* versus female *L. aldrichi*; Appendix Table A3), however, this metric was not a significant predictor in our multiple linear regression models. Due to this, we did not incorporate leg data into the remainder of the analyses. We found significant differences between species for all contact and association metrics, although the specific relationships varied. Overall, females and males of nonsacculate species (*L. vittatum* and *L. euserratipalpe*) associated significantly more closely than females and males of sacculate species (*L. bracchiolum* and *L. aldrichi*; Figure 2 & A2).

**Figure 2:**
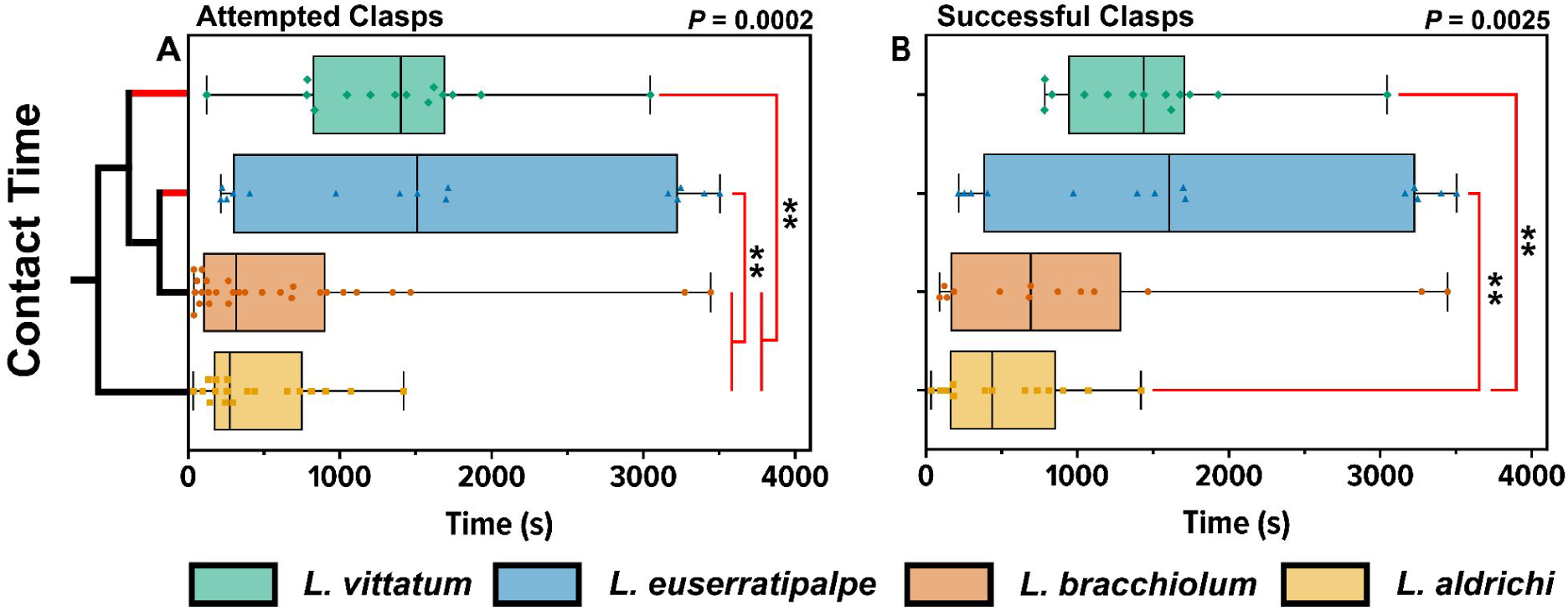
Association metrics between male and female pairs. Boxes represent medians plotted with IQR, symbols represent individual trial values, and whiskers correspond to the minimum and maximum trial values. The results from group comparison tests are shown in the upper right-hand corners, while significant results from post-hoc multiple comparisons tests are noted by red brackets with asterisks (* = *P* < 0.05; ** = *P <* 0.01). The simplified phylogeny indicates reproductive morphology; red tips indicate nonsacculate species and black tips represent sacculate species (phylogeny adapted from Burns et al. 2013). **A)** Total contact time in seconds between males and females during attempted-clasp trials. **B)** Total contact time in seconds between males and females during successful-clasp trials.

In attempted-clasp trials, nonsacculate males and females were in physical contact for significantly longer than sacculate males and females. In these trials, median contact time was between three and four times longer in nonsacculate species than in sacculate species (Figure 2A). When comparing only successful-clasp trials, both nonsacculate species retained significantly longer contact time when compared to *L. aldrichi*, however, *L. bracchiolum* was not significantly different (Figure 2B). When comparing the number of bouts of investigatory tapping performed during a mating trial, we found that *L. vittatum* did so significantly less than *L. bracchiolum* in both attempted-clasp (*P* = 0.001) and successful clasp trials (*P* = 0.005; Figure 3). Surprisingly, we also found that *L. vittatum* had significantly fewer bouts than its fellow nonsacculate in both trial categories (attempted-clasp trials: P = 0.0341; successful clasp trials: *P* = 0.0221), and that *L. bracchiolum* had significantly more tapping bouts than its fellow sacculate in attempted-clasp trials (*P* = 0.0208; Figure 3). There were no clear patterns when comparing flee instances between species, though *L. bracchiolum* females rarely fled males and had a median flee count of 0 instances (IQR 0 - 2 instances), resulting in L. bracchiolum having a significantly lower flee count than *L. aldrichi* in attempted-clasp (*P* > 0.0001) and successful clasp trials (*P* = 0.0121; Figure A3).

**Figure 3:**
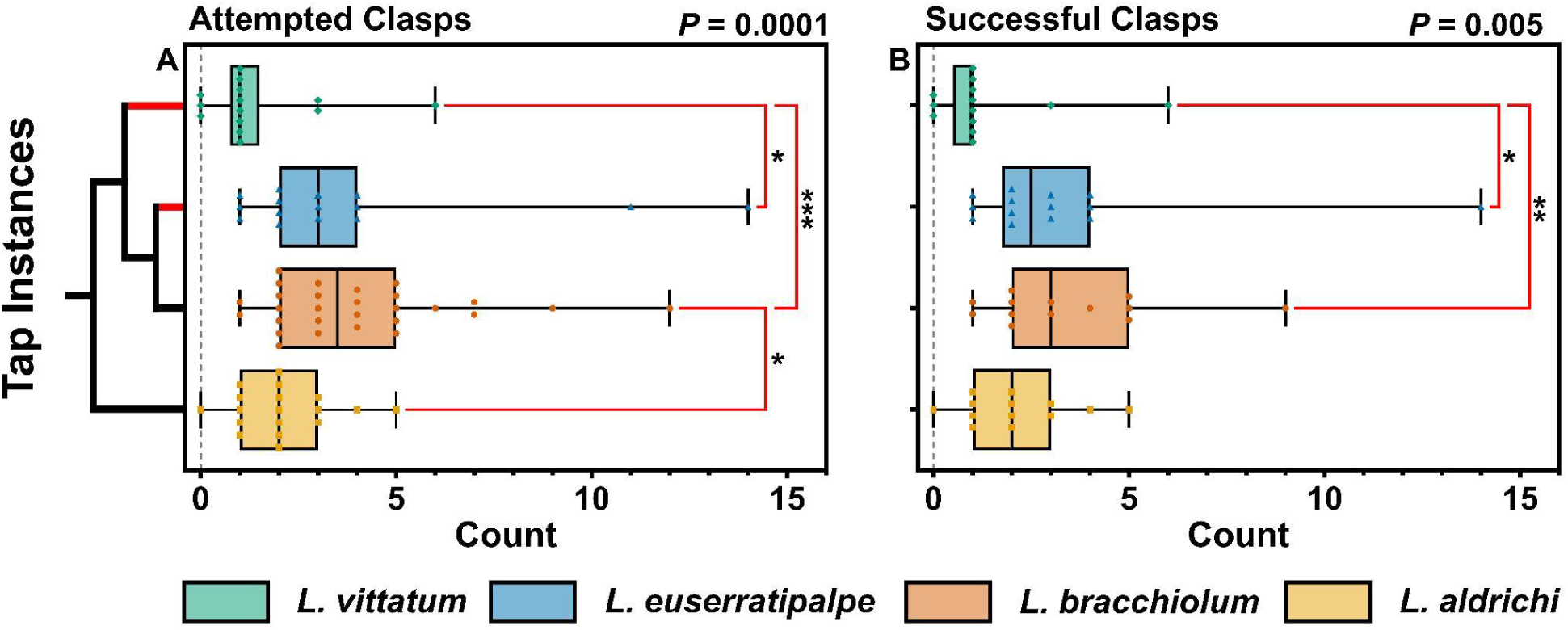
Instances of investigatory tapping bouts. Boxes represent medians plotted with IQR, symbols represent individual trial values, and whiskers correspond to the minimum and maximum trial values. The results from group comparison tests are shown in the upper right-hand corners, while significant results from post-hoc multiple comparisons tests are noted by red brackets with asterisks (* = *P* < 0.05; ** = *P* < 0.01; *** = *P* < 0.001). The simplified phylogeny indicates reproductive morphology; red tips indicate nonsacculate species and black tips represent sacculate species (phylogeny adapted from Burns et al. 2013). **A)** Instances of tapping bouts between males and females during attempted-clasp trials. **B)** Instances of tapping bouts between males and females during successful-clasp trials.

Using automated tracking, we found that in both attempted- and successful-clasp trials males were within one leg span of females significantly more often in nonsacculate species than in the nonsacculate *L. aldrichi*. This matched the contact patterns found in the successful-clasp trials manual scoring, however, for attempted-clasp trials we failed to find significance comparing *L. bracchiolum* to nonsacculates. This discrepancy in contact time is likely due to using the body centroids for position in TRex – we only considered females to be within the male’s leg span if the female’s body centroid was within the given male leg span, meaning leg-leg contact may occur without being scored. During attempted-clasp trials, nonsacculate females and males were also significantly closer to one another when compared to *L. aldrichi*, though this pattern did not hold for successful-clasp trials.

### Precopulatory Phase

Across all species, we found significant differences in the latency between first contact and first clasping attempt (Figure 4). *Leiobunum euserratipalpe* median attempt latency was 6.0 s (IQR 3.0 - 10.0 s), significantly lower than *L. vittatum* (76.5 s, IQR 20.75 - 605.3 s) and *L. aldrichi* (128.0 s, IQR 7.25 - 462.0 s), but was not significantly different from that of *L. bracchiolum* (18.0 s, IQR 4.0 - 327.5 s). All species displayed marked variation in latency to attempt a clasp except *L. euserratipalpe*, which had a standard deviation of 10.72 seconds (*L. bracchiolum* SD = 230.7 s; *L. aldrichi* SD = 363.5 s; *L. vittatum* SD = 330.4 s). The longest attempt latency for *L. euserratipalpe* was just 41 s, with all but three males making an attempt within 10 s of their initial contact with a female (Figure 4).

**Figure 4:**
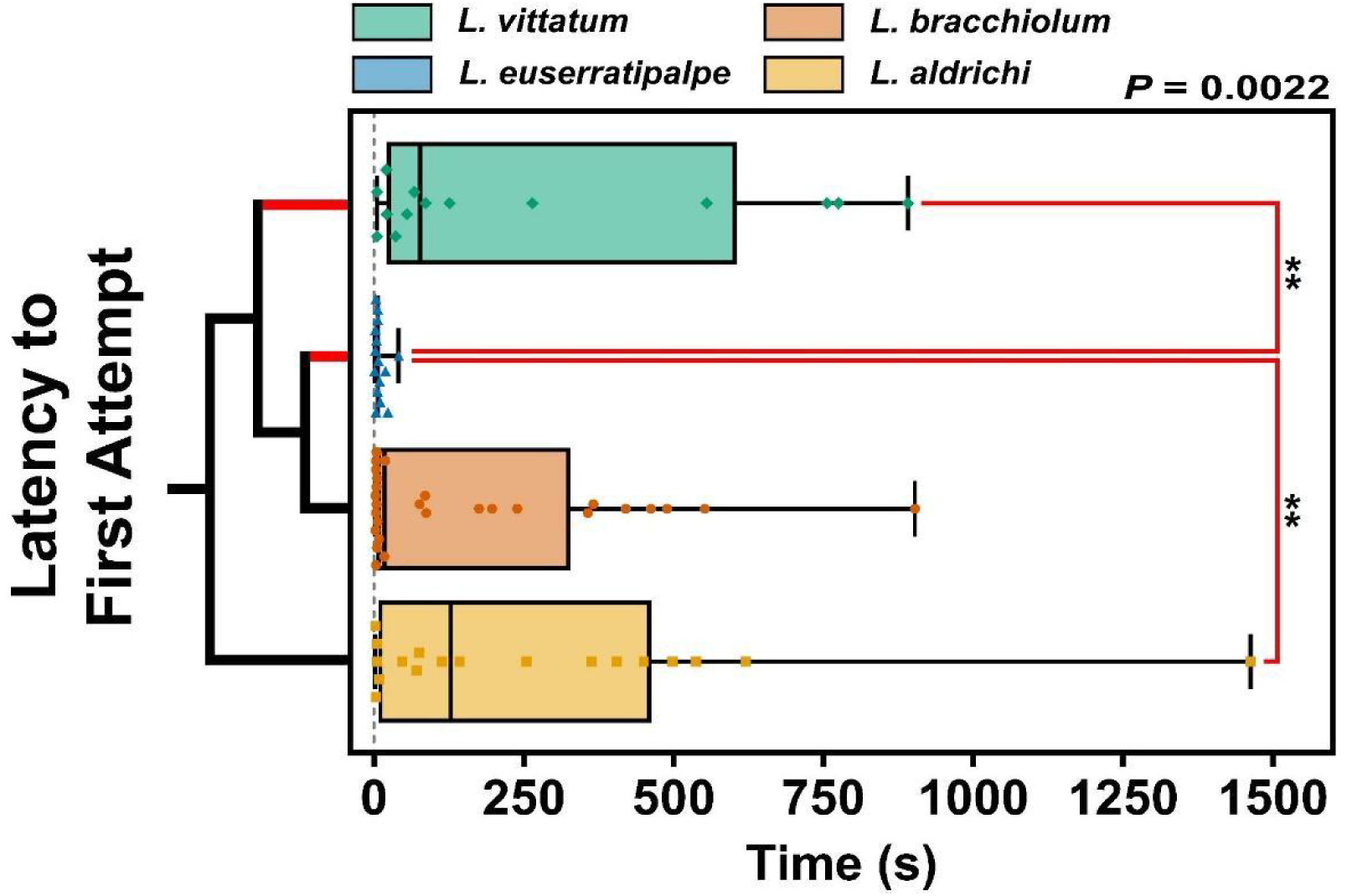
Latency in seconds between first contact and first clasp attempt during successful- clasp trials. Boxes represent medians plotted with IQR, symbols represent within-trial values, and whiskers correspond to the minimum and maximum trial values. The result from a Kruskal-Wallis test is shown in the upper right-hand corner, while significant results from Dunn’s multiple comparisons tests are noted by red brackets with asterisks (** = *P <* 0.01). The simplified phylogeny indicates reproductive morphology; red tips indicate nonsacculate species and black tips represent sacculate species (phylogeny adapted from Burns et al. 2013).

Finally, we found significant differences in pursuit angle between species for attempted- and successful-clasp trials. In trials with attempted clasps, the median pursuit angle for *L. vittatum* (82.64 degrees, IQR 82.05 - 85.45 degrees) was significantly smaller than that of *L. bracchiolum* (87.75 degrees, IQR 85.79 - 88.86 degrees) (Figure 5A). *Leiobunum vittatum* also had a significantly smaller angle than that of *L. euserratipalpe* (88.71 degrees, IQR 85.67 - 89.92 degrees) despite both being nonsacculate. Though in successful-clasp trials the *L. vittatum* pursuit angle (82.30 degrees, IQR 82.00 - 84.66 degrees) was similar to both sacculate species, it remained significantly smaller than *L. euserratipalpe* (89.04 degrees, IQR 85.53 - 89.89 degrees) (Figure 5B).

**Figure 5:**
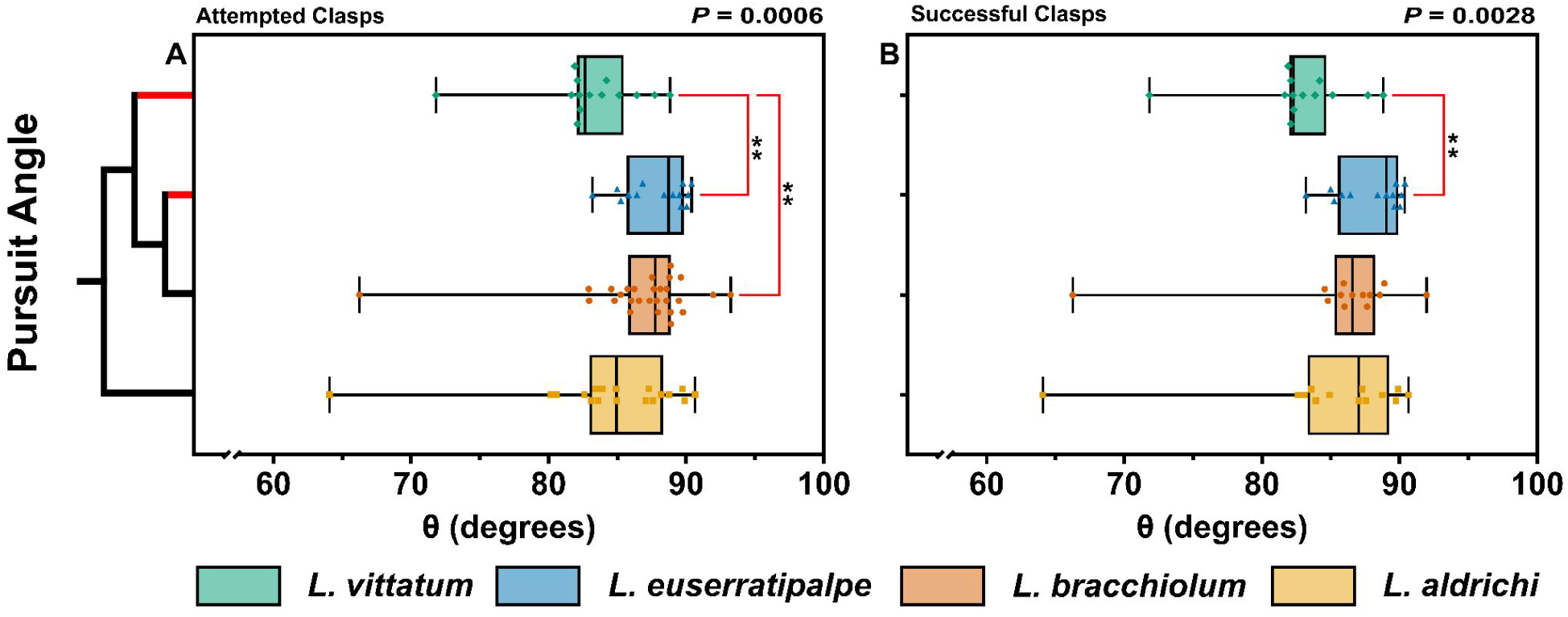
Angle between male and female trajectories between first contact and first clasp attempt. Boxes represent medians plotted with IQR, symbols represent within-trial means, and whiskers correspond to the minimum and maximum trial means. The simplified phylogeny indicates reproductive morphology; red tips indicate nonsacculate species and black tips represent sacculate species (phylogeny adapted from Burns et al. 2013). The results from Kruskal-Wallis tests are shown in the upper right-hand corner, while significant results from Dunn’s multiple comparisons tests are noted by red brackets with asterisks (** = *P <* 0.01). **A)** Attempted-clasp trials. **B)** Successful-clasp trials.

### Copulatory Phase

We found significant differences when comparing the overall clasping duration of each species using a Kruskal-Wallis test, although there were no clear patterns between sacculate and nonsacculate species (Figure 6). During successful-clasp trials *L. vittatum* males clasped females significantly longer than either sacculate species, with a median time of 676.8 s (IQR 310.6 - 1271.0 s) compared to 118.7 s (IQR 98.47 - 179.1 s) for *L. bracchiolum* and 260.5 s (IQR 17.11 - 662.8 s) for *L. aldrichi* (Figure 6). Interestingly it appeared that *L. euserratipalpe* clasped far less than expected, and an unplanned post-hoc test comparing the species to *L. vittatum* revealed that *L. euserratipalpe* clasped for significantly less time than its nonsacculate counterpart (Figure 6).

**Figure 6:**
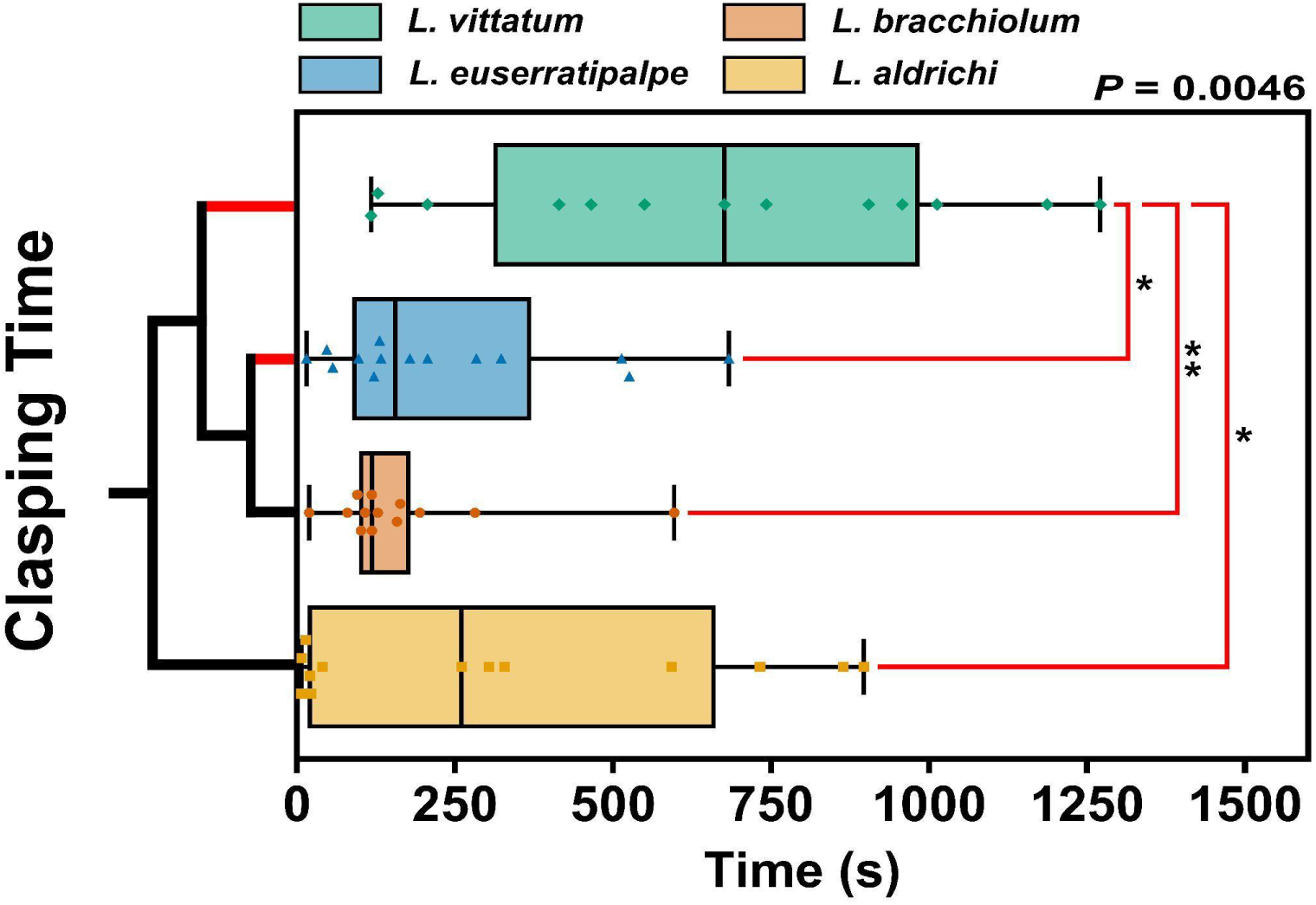
Time in seconds that males clasping females during trials with successful clasps. Boxes represent medians plotted with IQR, symbols represent within-trial values, and whiskers correspond to the minimum and maximum trial values. The result from a Kruskal-Wallis test is shown in the upper right-hand corner, while significant results from Dunn’s multiple comparisons tests are noted by red brackets with asterisks (* = *P* < 0.05; ** = *P <* 0.01). The simplified phylogeny indicates reproductive morphology; red tips indicate nonsacculate species and black tips represent sacculate species (phylogeny adapted from Burns et al. 2013).

The median clasping duration for *L. euserratipalpe*, 155.7 s (IQR 87.22 - 370.7 s), was more aligned with the clasping durations of sacculate species, and was approximately half that of *L. aldrichi*, contrary to our predictions.

Comparing the degree to which females angled their body (which blocks genital operculum access, Figure 1C) revealed significant differences across all species, but these analyses resulted in mixed results when comparing sacculate and nonsacculate species. *Leiobunum vittatum* was again significantly different from its nonsacculate counterpart, *L. euserratipalpe*, and the sacculate *L. aldrichi*, but it was not significantly different from *L. bracchiolum*. In all trials with attempted clasps, *L. vittatum* females angled their bodies to a median of 46.39 degrees (IQR 33.54 - 90.0 degrees) with four females reaching a full 90 degrees perpendicular to the substrate (Figure 7A).

**Figure 7:**
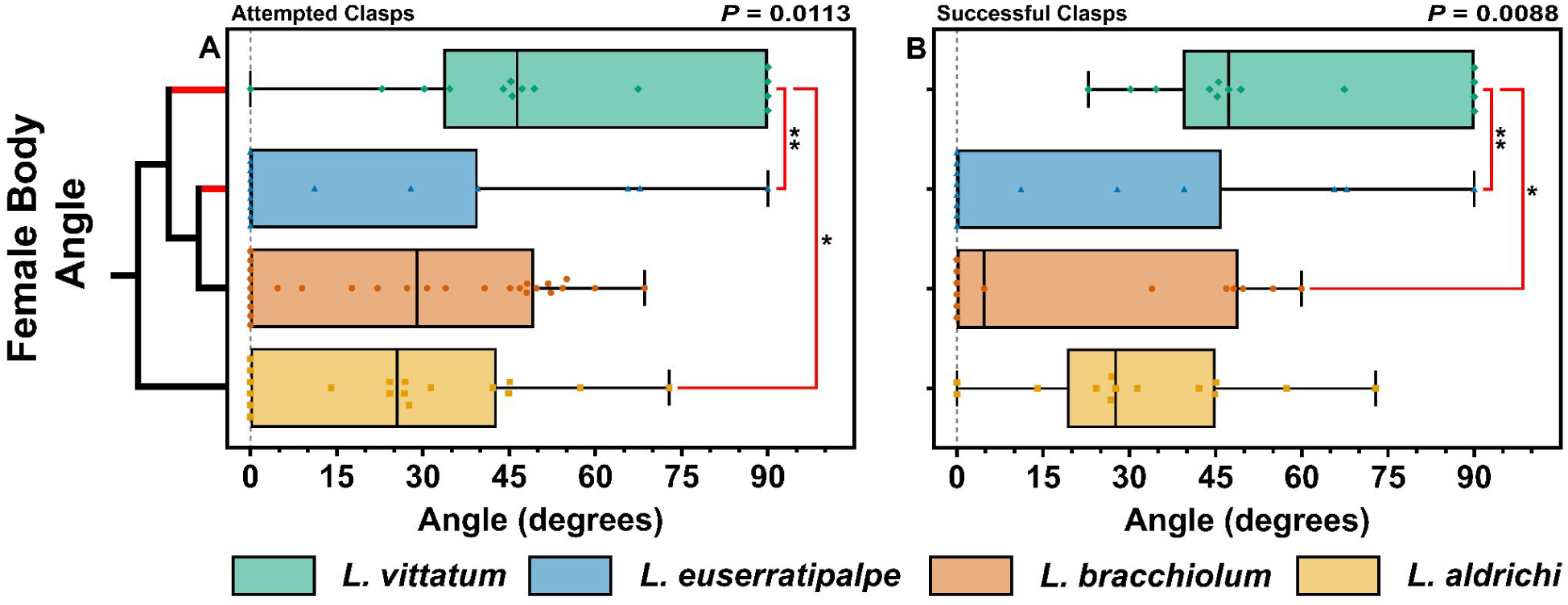
Maximum angle females formed between genital operculum and substrate. Boxes represent medians plotted with IQR, symbols represent within-trial values, and whiskers correspond to the minimum and maximum trial values. The simplified phylogeny indicates reproductive morphology; red tips indicate nonsacculate species and black tips represent sacculate species (phylogeny adapted from Burns et al. 2013). The results from Kruskal-Wallis tests are shown in the upper right-hand corner, while significant results from Dunn’s multiple comparisons tests are noted by red brackets with asterisks (* = *P* < 0.05; ** = *P <* 0.01). **A)** Attempted-clasp trials. **B)** Successful-clasp trials.

*Leiobunum euserratipalpe* females displayed a median angle of 0 degrees + (IQR 0 - 39.54 degrees) and rarely angled their bodies, with only six of 15 females doing so and only one female reaching 90 degrees. While *L. bracchiolum* and *L. aldrichi* females angled their bodies more often than *L. euserratipalpe* females, no females reached 90 degrees, and their median angles were 28.98 degrees (IQR 0 - 49.35 degrees) and 25.5 degrees + (IQR 0 - 42.84 degrees) respectively.

The contact time (between first touch and first clasp attempt) and the total time within one leg span were significant predictors of clasping success for both the total-species full model and the total-species reduced model (Table 3). Species-level models for *L. bracchiolum* and *L. aldrichi* were non-significant, while nonsacculate, *L. vittatum*, and *L. euserratipalpe* models were not possible due to few failed clasping attempts.

**Table 3:** Multiple logistic regression modelling of clasping success with behavioural metrics as predictor variables for the total-species full model and the total-species reduced model. Predictor variables were iteratively removed beginning with the variable with the greatest *P*-value until no variables with *P* > 0.1 remained. Significant predictor variables are in bold.

| **Model/Factor** | **AICc** | **Pseudo-R<sup>2</sup>** | **Estimate** | **SE** | **\|Z\|** | **P-value** |
| --- | --- | --- | --- | --- | --- | --- |
| *Total-species full model* | 72.95 | 0.388 | - | - | - | - |
| Intercept | - | - | -3.247 | 8.348 | 0.389 | 0.6973 |
| Initial latency | - | - | -0.002 | 0.002 | 1.125 | 0.2607 |
| **Pre-attempt contact time** | - | - | **-0.004** | **0.002** | **2.482** | **0.0131** |
| Average distance | - | - | 0.114 | 0.093 | 1.226 | 0.2200 |
| **Time within 1 leg span** | - | - | **0.003** | **0.001** | **2.257** | **0.0240** |
| Female body angle | - | - | 0.032 | 0.017 | 1.885 | 0.0594 |
| Pursuit angle | - | - | 0.016 | 0.095 | 0.168 | 0.8665 |
| Flee count | - | - | 0.606 | 0.438 | 1.385 | 0.1661 |
| Tap count | - | - | -0.216 | 0.125 | 1.721 | 0.0853 |
| *Total-species reduced model* | 68.87 | 0.319 | - | - | - | - |
| Intercept | - | - | -1.113 | 0.645 | 1.725 | 0.0845 |
| **Pre-attempt contact time** | - | - | **-0.003** | **0.001** | **2.685** | **0.0072** |
| **Time within 1 leg span** | - | - | **0.002** | **7e-04** | **2.844** | **0.0045** |
| Female body angle | - | - | 0.024 | 0.014 | 1.690 | 0.0910 |
| Flee count | - | - | 0.755 | 0.391 | 1.928 | 0.0538 |

### Postcopulatory

Finally, when analysing the duration of mate guarding – the primary indicator of postcopulatory conflict in leiobunine harvesters – we found significant differences across species (Figure 8). Nonetheless, there were few pairwise significant differences, potentially due to significant intraspecific variation in all species except *L. aldrichi*. *Leiobunum aldrichi* males guarded for a median time of just 59.0 s + (IQR 0 - 270.5 s), while *L. bracchiolum* and *L. vittatum* males guarded for 474.0 s (IQR 0 - 857.0 s) and 351 s (IQR 18.0 - 1145.0 s), respectively (Figure 8). *Leiobunum euserratipalpe* males guarded females for significantly longer than *L. aldrichi* males did, with its median guarding times being 1209.0 s (IQR 209.8 - 2919.0 s) (Figure 8). Although male *L. euserratipalpe* tended to guard females for extended periods, they did not always do so – five males guarded for less than 320.0 s, and two of these did not guard at all. Due to its broad distribution of guarding duration, we did not have sufficient statistical power to resolve any potential differences between *L. euserratipalpe*, *L. vittatum*, and *L. bracchiolum*, and further study may allow the resolution of these differences.

**Figure 8:**
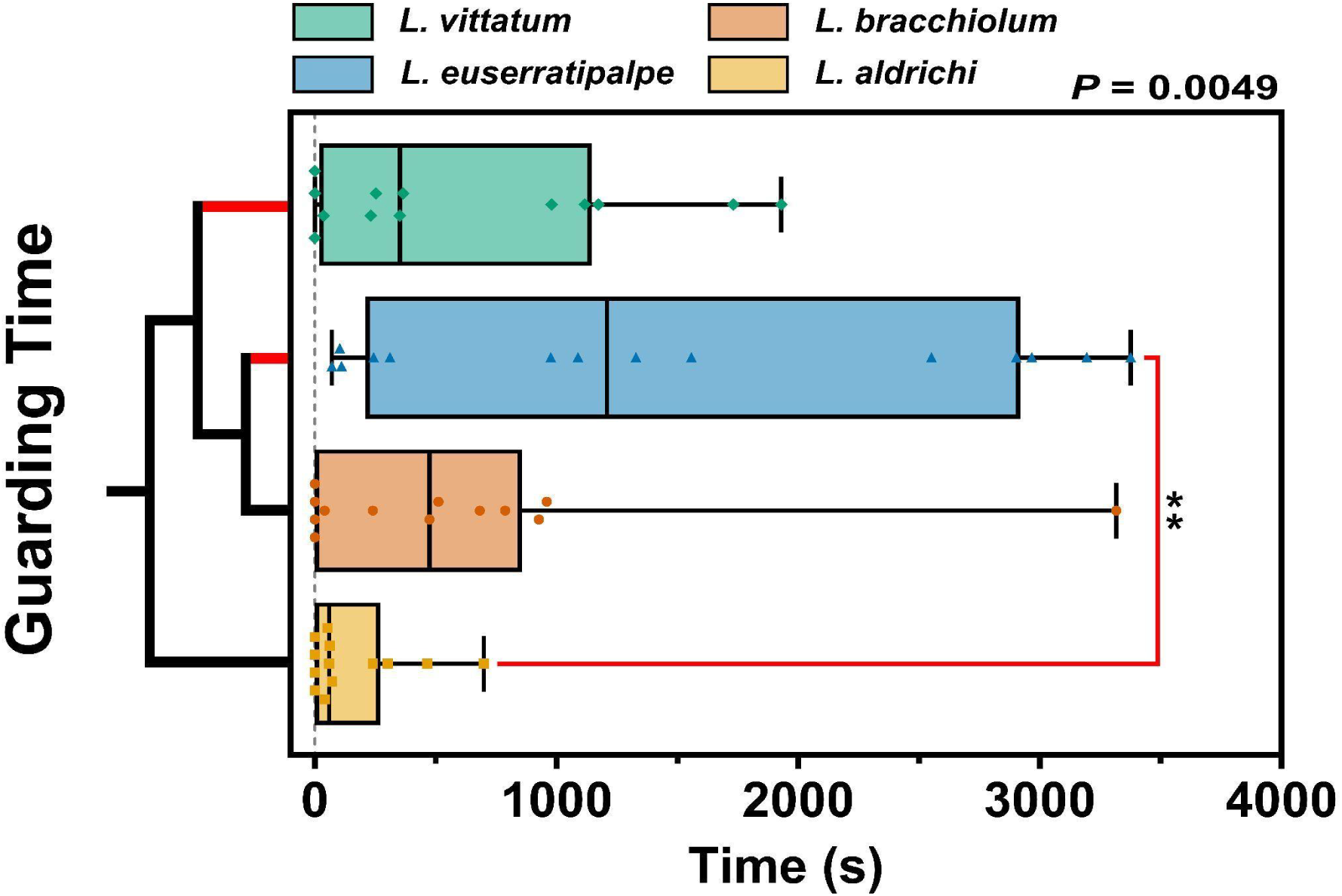
Time in seconds that males guarded females during successful-clasp trials. Boxes represent medians plotted with IQR, symbols represent within-trial values, and whiskers correspond to the minimum and maximum trial values. The result from a Kruskal-Wallis test is shown in the upper right-hand corner, while significant results from Dunn’s multiple comparisons tests are noted by red brackets with asterisks (** = *P <* 0.01). The simplified phylogeny indicates reproductive morphology; red tips indicate nonsacculate species and black tips represent sacculate species (phylogeny adapted from Burns et al. 2013).

## Discussion

### Behavioural comparisons between sacculate and nonsacculate species

Our results indicate that the nonsacculate harvesters *L. vittatum* and *L. euserratipalpe* display significantly different mating behaviour when compared to the sacculate species *L. aldrichi* and *L. bracchiolum* (Table 4). This variation in mating behaviour corresponds to the penis morphology and reproductive syndromes of each species; *L. vittatum* and *L. euserratipalpe* lack precopulatory nuptial gifts and possess antagonistic morphology, while *L. aldrichi* and *L. bracchiolum* give precopulatory nuptial gifts and lack antagonistic morphology (Burns et al. 2013). Across all mating trials, nonsacculate males repeatedly displayed significantly longer median association times, were significantly closer to females, and performed fewer investigatory tapping bouts (Figures 2, 3, & A2). Male *L. euserratipalpe* attempted mating clasps significantly faster upon first touch than male *L. aldrichi* and *L. vittatum* (Figure 4), while male *L. vittatum* pursued females more vigorously and female *L. vittatum* also displayed higher levels of resistance behaviour (Figures 5, 7, & A3). Hallmark conflict behaviours such as extended mate clasping and mate guarding, were not found to be elevated across both nonsacculate species, with instead *L. vittatum* displaying extended clasping behaviour (Figure 6) and *L. euserratipalpe* displaying extended guarding behaviour (Figure 8). Altogether, these data indicate that nonsacculate harvesters display significantly higher levels of antagonistic behaviour, possibly to compensate for the loss of precopulatory nuptial gifts and the reduction of pericopulatory nuptial gifts (Table 4).

**Table 4:** Summary of our predictions for behavioural sexual conflict intensity and the degree to which the results of each metric matched what we expected for each species (see Table 2 for metric specific expectations). “Support” indicates that the species matched our predictions in both attempted-clasp and successful-clasp trials; “Mixed” indicates metrics in which this occurred in only one of the two trial categories (attempted-clasp and successful-clasp), and “No support” indicates that the species did not match our expectations in either trial category.

| Metric | *L. vittatum* | *L. euserratipalpe* | *L. bracchiolum* | *L. aldrichi* |
| --- | --- | --- | --- | --- |
| Conflict prediction | High | High | Low | Low |
| Latency to 1st attempt | No Support | Support | No Support | Support |
| Pursuit angle | Support | No Support | Mixed | No Support |
| Contact time | Support | Support | Mixed | Support |
| Tap Count | Support | No Support | Support | No Support |
| Flee Count | No Support | Mixed | Mixed | No Support |
| Time within 1 leg span | Support | Support | No Support | Support |
| Pairwise distance | Mixed | Mixed | No Support | Mixed |
| Female body angle | Support | No Support | Mixed | Mixed |
| Clasping time | Support | No Support | Support | Support |
| Guarding time | No Support | Support | No Support | Support |
| Supporting Metrics | 6-7 of 10 | 4-6 of 10 | 2-6 of 10 | 5-7 of 10 |

With these results we provide the first direct behavioural comparison between sacculate and nonsacculate leiobunine harvesters. Despite extensive evidence of a shift from a solicitous suite of traits to an antagonistic suite, the role of behaviour had not yet been investigated with respect to sexual conflict in this system. Previous studies have revealed various losses of solicitous traits such as penile sacs, precopulatory nuptial gifts, and high essential amino acid content in pericopulatory nuptial gifts (Burns et al. 2013; Kahn et al. 2018). Meanwhile, these and other studies have also shown gains of antagonistic traits, such as enhanced penile prying force, female reproductive barriers, and modified clasping structures (Burns et al. 2013; Burns & Shultz 2015, 2016; Karachiwalla et al. 2020). While these changes arose independently across leiobunine harvesters, many have reached convergent states. Our results indicate that these morphological and biochemical changes have been accompanied by marked behavioural changes as well.

Due to leiobunine harvesters primarily relying on their antenniform legs for sensory cues (Shultz & Pinto-da-Rocha 2007; Willemart & Hebets 2012) we expected that males would keep females within their leg span throughout antagonistic mating interactions to facilitate control. In this study, nonsacculate species maintained this state for two to three times longer than sacculate species. The distance and contact time between individuals are not clear indicators of sexual conflict in isolation – both can be expected to occur in solicitous and antagonistic interactions (though short contact time would primarily indicate a lack of antagonism due to extended clasping and guarding being hallmark characteristics of behavioural conflict; Arnqvist & Rowe 2005; Bokides et al. 2012; Iglesias-Carrasco et al. 2019). However, in conjunction with the other metrics in the study, including female body angle, clasping time, and guarding time, it seems more likely to be associated with antagonism. Across mammals, reptiles, fish, and arthropods, many studies have shown cohabitation with males or proximity to males can each impact female fitness negatively through harassment (Arnqvist & Rowe 2005; Krupa et al. 1990; Kunz et al. 2021; Pilastro et al. 2003; Plath et al. 2003; Rönn et al. 2006; Schlupp et al. 2001; Schütz & Taborsky 2005; Shine et al. 2005). In particular, even if copulation does not occur, close association with males can lower female fitness, whether due to direct stress from male activity or from selecting low-quality refuge instead of a high-quality habitat with males as an avoidance strategy (Bokides et al. 2012; Iglesias-Carrasco et al. 2019). Furthermore, the exceptionally low latency between first contact and first attempt in *L. euserratipalpe* might be driven by a variety of factors, although many which might normally influence latency are controlled in this experiment, such as female age or sexual history, male starvation, and the presence of male rivals (Burke & Bonduriansky 2018; Dmitriew & Blanckenhorn 2012; Fowler et al. 2022). Low latency has previously been associated with high conflict (Blanckenhorn et al. 2020; Skwierzyńska & Plesnar-Bielak 2018), and in concert with the elevated guarding duration in *L. euserratipalpe,* this low latency appears to indicate a species with high conflict.

### Clasping, guarding, and conflict

Extended clasping time (beyond what might be ideal for female fitness) is a classic indicator of sexual conflict found in myriad arthropod and vertebrate systems (Arnqvist & Rowe 2005; Johns et al. 2009; Rowe & Arnqvist 2012; Rueda-Solano et al. 2022; Sigvardt et al. 2017). Indeed, males of some species, including those in this study, are entirely unable to ensure copulation when unable to clasp the female, providing significant motivation for unreceptive females to prevent clasping (Machado & Macías-Ordόñez et al. 2007; Myers et al. 2016). Extended clasping can function as a multifaceted advantage for males, the primary explanatory hypotheses being the Ejaculate Transfer (ET) hypothesis (Edvardsson & Canal 2006; Mazzi et al. 2009; Simmons 2002) and the Extended Mate Guarding (EMG) hypothesis (Alcock 1994; Mazzi et al. 2009). In the ET hypothesis, males extend clasping time to maximise the removal of sperm from previous males and the transfer of ejaculate materials such as sperm, seminal fluids, and mating plugs (Arnqvist & Rowe 2005; Edward et al. 2015; Mazzi et al. 2009; Macías-Ordόñez et al. 2010; Simmons 2002; Townsend et al. 2019; Vahed et al. 2014). In the EMG hypothesis, males instead maximise clasping time as a function of mate guarding in order to reduce the potential for female remating (Mazzi et al. 2009; Zatz et al. 2011). In both these nonexclusive hypotheses, the ultimate result may be quite similar to those achieved by nuptial gifts (Lewis & South 2012). Mating plugs, sperm removal, and clasping as mate guarding have all been observed in tropical harvesters (Machado & Burns 2023; Macías-Ordόñez et al. 2010; Townsend et al. 2015, 2019; Zatz et al. 2011), and so the behavioural conflict over clasping time in these North American harvesters might be attributable to either hypothesis or some combination of the two.

Female harvesters often angle their body anterior-down into the substrate, blocking access to their genital operculum. This potentially makes it more difficult for males to achieve or maintain a clasp, though this does not seem to be particularly effective for females in our study based on our logistic regression models (Figure 1C, Table 3; Fowler-Finn et al. 2014, 2018, 2019; Machado & Macías-Ordόñez 2007). If females orient their bodies in this manner, males cannot access the genital operculum and therefore cannot initiate intromission (Figure 1C). Many arthropod species have evolved morphological anti-clasping adaptations (Arnqvist & Rowe 1995, 2005; Karlsson Green et al. 2013) similarly, body angling might be considered a concordant behavioural anti-clasping adaptation – actions to move the genitals away from males are common among arthropods with high levels of sexual conflict (Akinyemi et al. 2021; Burke et al. 2015; Golov et al. 2018; Machado & Macías-Ordόñez 2007; Maroni et al. 2023; Zweerus et al. 2021). While our regression modelling did not include female body angle as a significant predictor, it is possible that the enhanced clasping morphology found in male *L. vittatum* allows them to effectively overcome this female behaviour, and further investigation is required.

Alternatively, there is some evidence that extended clasping time in *L. vittatum* may be attributable to female antagonism rather than male, which would also serve to explain the discrepancy between clasping times in the two nonsacculate species. First suggested by Fowler-Finn et al. (2014), female *L. vittatum* may clasp the male’s penis using the genital operculum, allowing the female to receive more nuptial gift than is optimal for the male to provide. Once clasping has been achieved and intromission has begun, male *L. vittatum* occasionally appear to struggle to end copulation (Fowler-Finn et al. 2014). This phenomenon is supported by genital morphology; female *L. vittatum* have well-developed opercular musculature and sclerotization, in stark contrast to the male’s relatively fragile penis (Burns & Shultz 2016). Burns and Shultz (2016) additionally cite reports of males with broken penises in the field, an injury consistent with struggles to disengage females. Female aggression can negatively correlate with nuptial gift quality and quantity (Kuriwada & Kasuya 2012; Toft & Albo 2016), and in *L. vittatum* this manipulation by females might be the direct result of the deterioration of nuptial gift quality and quantity (Khan et al. 2018; Machado & Burns 2023; Machado & Macías-Ordόñez 2007). Because females of nonsacculate species can only receive nuptial gift fluid during intromission, rather than both before and during, there may be a tangible benefit to females remaining in copula as long as possible. It may be that this acts as a cost-mitigating strategy for females, although it seems unlikely that females would prefer this as opposed to simply avoiding the encounter altogether due to the elevated levels of precopulatory resistance from females. The potential transition from male pre-copulatory antagonism to female copulatory antagonism would provide a fascinating dichotomy of sexual conflict deserving further research.

Mate guarding is another strong indicator of potential sexual conflict commonly found in arthropods (Alcock 1994; Arnqvist & Rowe 2005; Simmons 2002). Mate guarding may allow males additional control over female mating decisions by discouraging remating, preventing sperm removal by rival males, and reducing sperm competition – all factors which might alternatively (or additionally) be influenced by nuptial gifts (Alcock 1994; Arnqvist & Rowe 2005; Elias et al. 2014; Frankino & Sakaluk 1994; Gór et al. 2023; Hagg et al. 2024; Lewis & South 2012; Machado & Burns 2023; Sakaluk 1991; Simmons 2002). It is important to note, however, that mate guarding can be beneficial if the guarding male reduces future harassment or predation for the female, though in these species female resistance to guarding should be less prevalent (Rodríguez-Muñoz et al. 2011; Sherman 1983). Sperm competition and cryptic female choice are distinct possibilities in leiobunine harvesters – all females have a sperm-storing spermatheca, while some species have a complex, multi-chambered spermatheca potentially affording the female control over sperm fate (Karachiwalla et al. 2020; Shultz & Pinto-da-Rocha 2007). These potential confounding factors would only further incentivize males to guard females and prevent remating, as suggested by both models and experimental evidence (Bateman et al. 2001; Del Matto et al. 2021; Gór et al. 2023; Harts & Kokko 2013). Many male Opiliones, including *Leiobunum* species, engage in extended mate guarding which typically lasts until oviposition (Fowler-Finn et al. 2014, 2018, 2019; Machado & Burns 2023, Macías-Ordóñez 1997; Wijnhoven 2011; Zatz et al. 2011). This is likely an extremely costly behaviour for males because these harvesters often aggregate in high density, leading to elevated levels of male-male competition (Macías-Ordóñez 1997). The few field studies of *Leiobunum* mating behaviour have observed male *Leiobunum spp.* defending ovipositing females from upwards of 6 rival males, although evidence for late-season male sex-ratio biases and the dearth of field studies likely leads to this being an underestimate (Bacon, R. & Brown, T. A., pers. observ; Macías-Ordóñez 1997; Wijnhoven 2011). Our study did not find *L. vittatum* guarding time to be significantly longer than sacculate species, but a key missing factor may be the absence of competitors. In previous studies, male *L. vittatum* were found to be most aggressive when in the presence of both males and females during their reproductive period (Macías-Ordóñez 1997). While this aggression was often male-directed, it remains possible that the absence of male competitors resulted in less guarding than expected.

### Multiple mating strategies

The *calcar* species-group and *vittatum* species-group have convergently evolved similar nuptial gift chemistries and penis morphologies (Burns et al. 2013; Kahn et al. 2018), yet the groups vary significantly in sexual and somatic morphology and biomechanics (Burns & Shultz (2015, 2016). Female genital operculum morphology differs in reproductive barrier structure between groups as well (Burns et al. 2013; Ingianni et al. 2011). Despite females of both groups possessing unique, species-specific pregenital barriers formed through sclerotization of the operculum and sternum, the mechanism through which these structures engage varies between groups (Burns et al. 2013; Shultz 2018). *Leiobunum euserratipalpe* does not have sexually dimorphic palpal morphology, while *L. vittatum* does (Ingianni et al. 2011; Shultz 2018). This variation in somatic morphology correlates with the significant differences in clasping duration revealed in this study – *L. vittatum* males clasp longer than *L. euserratipalpe* and possess clasping structures indicative of this. *Leiobunum euserratipalpe* males, lacking enhanced palpal clasping structures, do not have particularly long clasping durations and instead guard females for extended periods postcopulation.

These variable suites of antagonistic traits indicate the evolutionary forces driving the transition to antagonism may differ between the *calcar* and *vittatum* species-groups, and further research investigating these mechanisms is needed. There are many temperate species within the Opiliones family Sclerosomatidae with documented transitions towards nonsacculate morphology, but to date no tropical species with these transitions have been reported (Burns et al. 2013; Burns & Tsurusaki 2016; Machado & Burns 2023; Martens 2020). While there is a dearth of tropical sexual conflict research overall, neotropical Opiliones have been fairly well studied, albeit with a focus on non-Sclerosomatidae species (Bacon et al. 2023; Burns & Machado 2023; Pinto-da-Rocha et al. 2007). This conspicuous lack of ostensibly antagonistic structures in tropical Sclerosomatidae might indicate that harsher climes drive the transition towards antagonism in temperate sclerosomatids. These data, in conjunction with previous morphological, phylogenetic, and biochemical work (Burns et al. 2013; Burns & Shultz 2015, 2016; Kahn et al. 2018), suggest nonsacculate *Leiobunum* harvesters have a frequently coercive mating system driven by sexual conflict, but this is not to say that every mating interaction is inherently antagonistic.

An alternative explanation for the behaviour we observed is cryptic female choice (CFC), which has been proposed to explain many common morphological and behavioural patterns typically assumed to be due to sexually antagonistic coevolution (Eberhard 1996, 2010). Under CFC, females are expected to exhibit peri- or post-copulatory preferences for particular male genital morphologies, such as those better able to stimulate the female (Eberhard 1996, 2010). The female may then preferentially store sperm, fertilise eggs, or reduce its remating rate to the benefit of the male (Eberhard 1996, 2010). Eberhard (2011) presents an excellent framework for effectively testing whether male stimulation of the female occurs by experimentally “blinding” a female to potential stimulation. Myers et al. (2016) performed such an experiment on the stick insect *Clitarchus hookeri*, which exhibits mating behaviour similar to Opiliones in that males must clasp females for mating to occur, and females have a moveable genital operculum. Myers et al. (2016) demonstrate through female desensitisation and male clasper abrasion that this clasping is not antagonistic and likely serves to signal to females to open their genital operculum and allow intromission. Though they note that, unlike in the present study, unmodified females typically do not resist unmodified males, their study nonetheless presents an excellent example of necessary future work in *Leiobunum*.

## Conclusions

While we intended to expedite video analysis of reproductive behaviour through a combination of automated tracking and manual scoring, we found the benefits of incorporating automated tracking to be minimal. Our subsequent attempts using alternative tracking programs resulted in similar difficulties, and many fine-scale behaviours, such as those involving leg or pedipalp movements, were not feasible to score with automated tracking and required manual scoring. Automated tracking may still provide benefits when used across many studies because trained models can be reused within a species, but manual tracking likely remains preferable for species with fine-scale behaviour or rapid positional changes, or when de novo training is required.

Our study demonstrates that mating behaviour significantly differs between sacculate and nonsacculate species, providing another line of evidence that sexual conflict has shaped reproduction in these Opiliones. Our data suggests that nonsacculate harvesters may resort to antagonistic behaviours in lieu of nuptial gifts to increase fitness through some combination of copulation success, insemination success, sperm transfer, sperm storage, and paternity share. Importantly, genital modification studies and economic studies of the costs of mating remain to be done in *Leiobunum* harvesters. Though it remains possible that other mechanisms, such as cryptic female choice, play an important role in this genus, it is unlikely for sexual conflict to be entirely absent in a system in which there is benefit for males to circumvent female preference (Snow et al. 2019).

A significant area that remains unclear is the effect of reproductive phenology on sexual conflict. North American leiobunine harvesters have significant variation in the timing of major life events such as maturation, oviposition, and senescence, as well as in the length of the reproductive period (Curtis & Machado 2007; Machado & Macías-Ordόñez 2007). Additionally, male-male competition and female antagonism are further factors which likely contribute to the evolution and maintenance of male antagonism in leiobunine Opiliones. How these myriad factors interact and impact sexual conflict is not well established, and the evidence of significant variation in mating behaviour in leiobunine harvesters provides the perfect opportunity for further behavioural research.

## Author Contributions

**Tyler A. Brown**: Conceptualization, methodology, investigation, formal analysis, visualisation, writing – original draft, writing – review & editing, funding acquisition. **Emily Marinko**: Investigation, formal analysis, writing – review & editing. **Mercedes Burns**: Conceptualization, methodology, formal analysis, writing – review & editing, supervision, funding acquisition.

## Data Availability

Mendeley Data doi: https://doi.org/10.17632/gsvd5z9g2b.2

## Acknowledgements

We would like to thank Ryan Bacon, Sai Wai, Emily Redmond, Lian Jackson, and Shelby Hawk for assistance with specimen collection and husbandry. We would also like to thank Sophia Nawaz and Megan Ramirez Cuenca for data processing assistance and Dr. Andrew Gordus for his helpful advice regarding automated tracking implementation. T. A. B. and M. B. were supported by funds from NSF IOS 00116. This work was also funded in part by the American Arachnological Society and Washington Biologists’ Field Club, which also provided access to Plummer’s Island.

## Appendix

**Figure A1:**
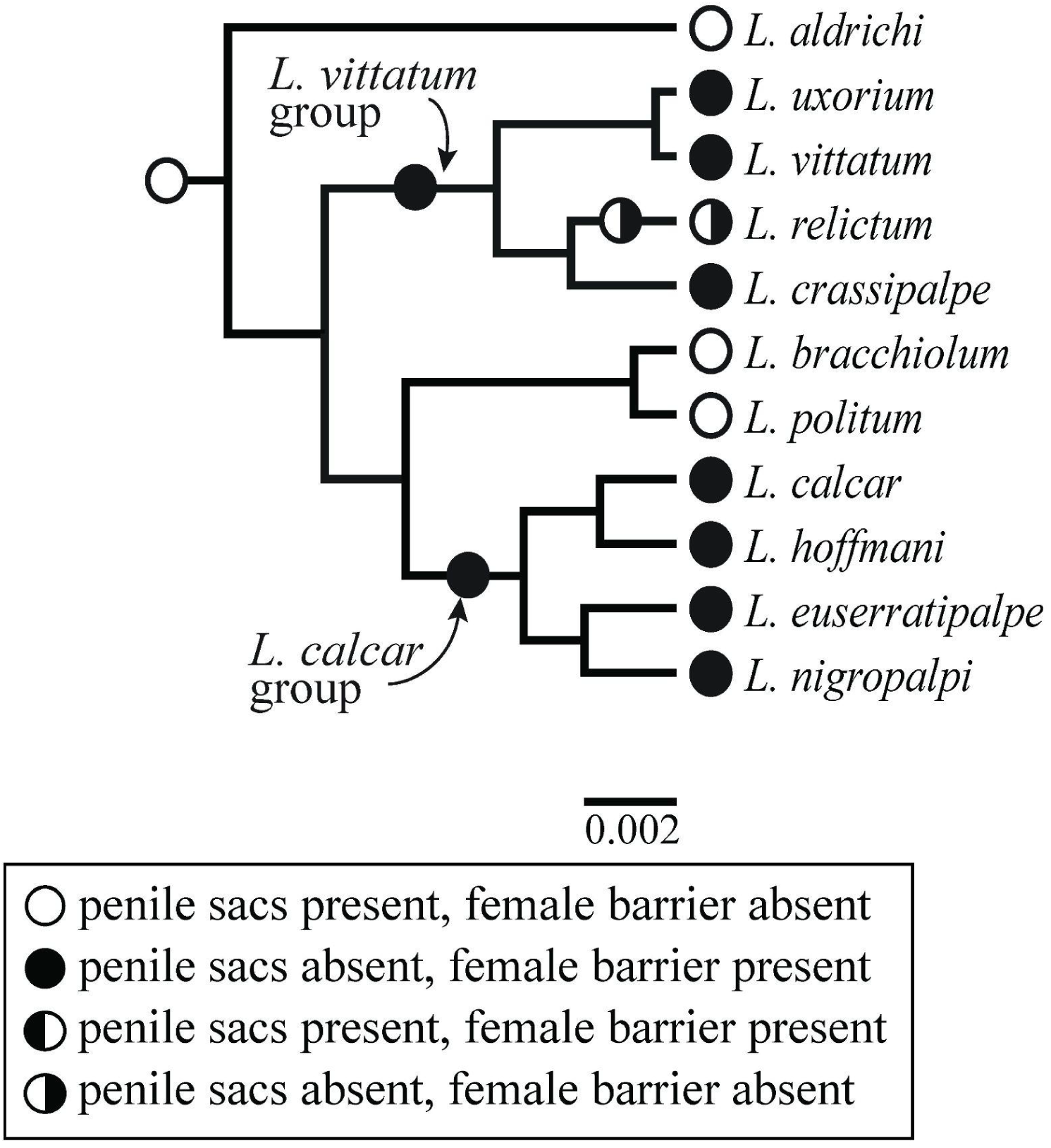
Bayesian Maximum Clade Credibility tree of the *Leiobunum* species used in this study, adapted from Burns & Shultz (2015), where scale = base substitutions/site. Parsimonious character state combinations of penile sac presence and female pregenital barrier presence are mapped on branches.

**Figure A2:**
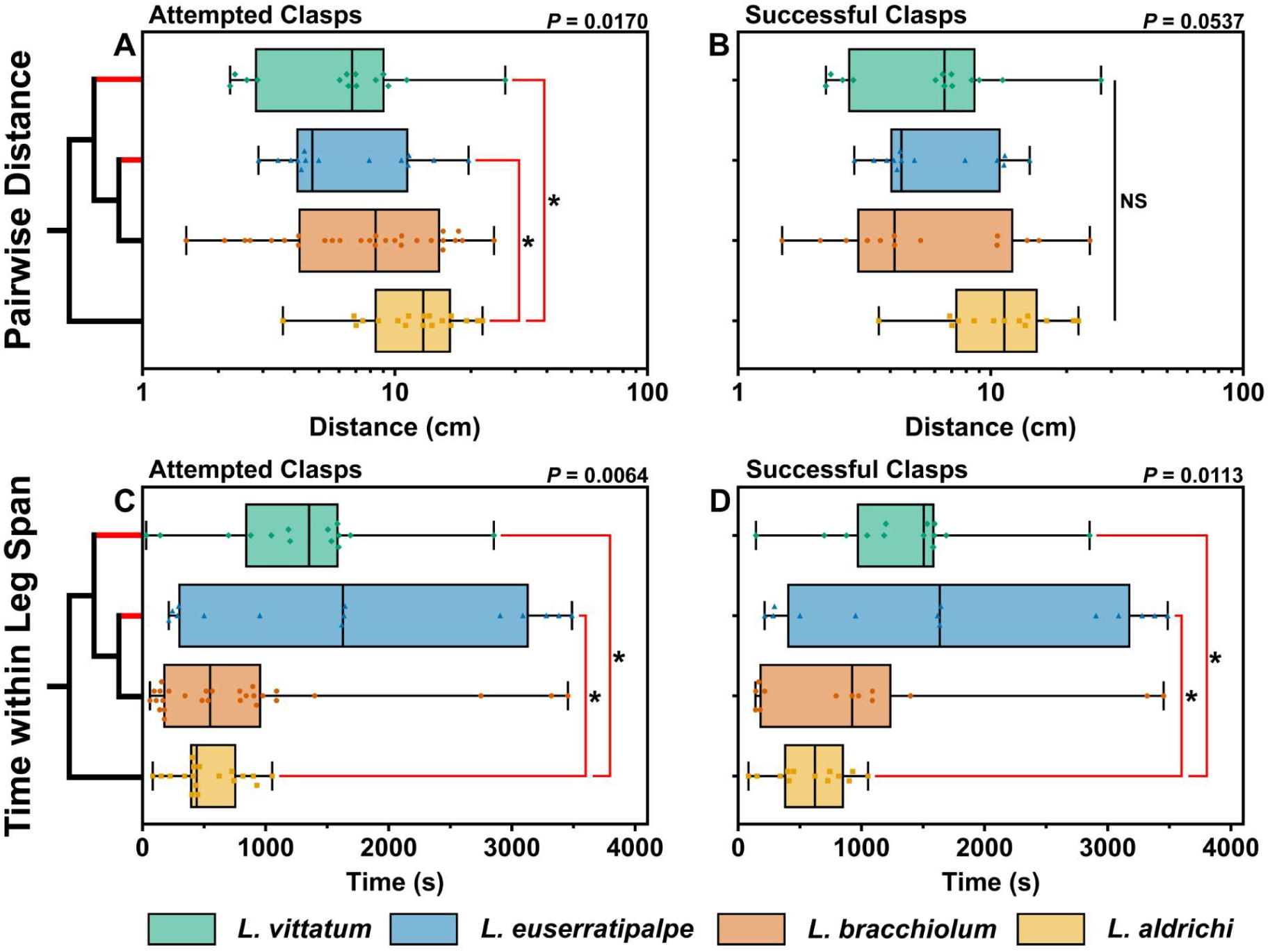
Association metrics between male and female pairs. Boxes represent medians plotted with IQR, symbols represent individual trial values, and whiskers correspond to the minimum and maximum trial values. The results from group comparison tests are shown in the upper right-hand corners, while significant results from post-hoc multiple comparisons tests are noted by red brackets with asterisks (* = *P* < 0.05). The simplified phylogeny indicates reproductive morphology; red tips indicate nonsacculate species and black tips represent sacculate species (phylogeny adapted from Burns et al. 2013). **A)** Average distance in centimetres between male and female pairs during attempted-clasp trials. **B)** Average distance in centimetres between male and female pairs during successful-clasp trials. **C)** Total time in seconds the female was within one leg span of the male during attempted-clasp trials. **D)** Total time in seconds the female was within one leg span of the male during successful-clasp trials.

**Figure A3:**
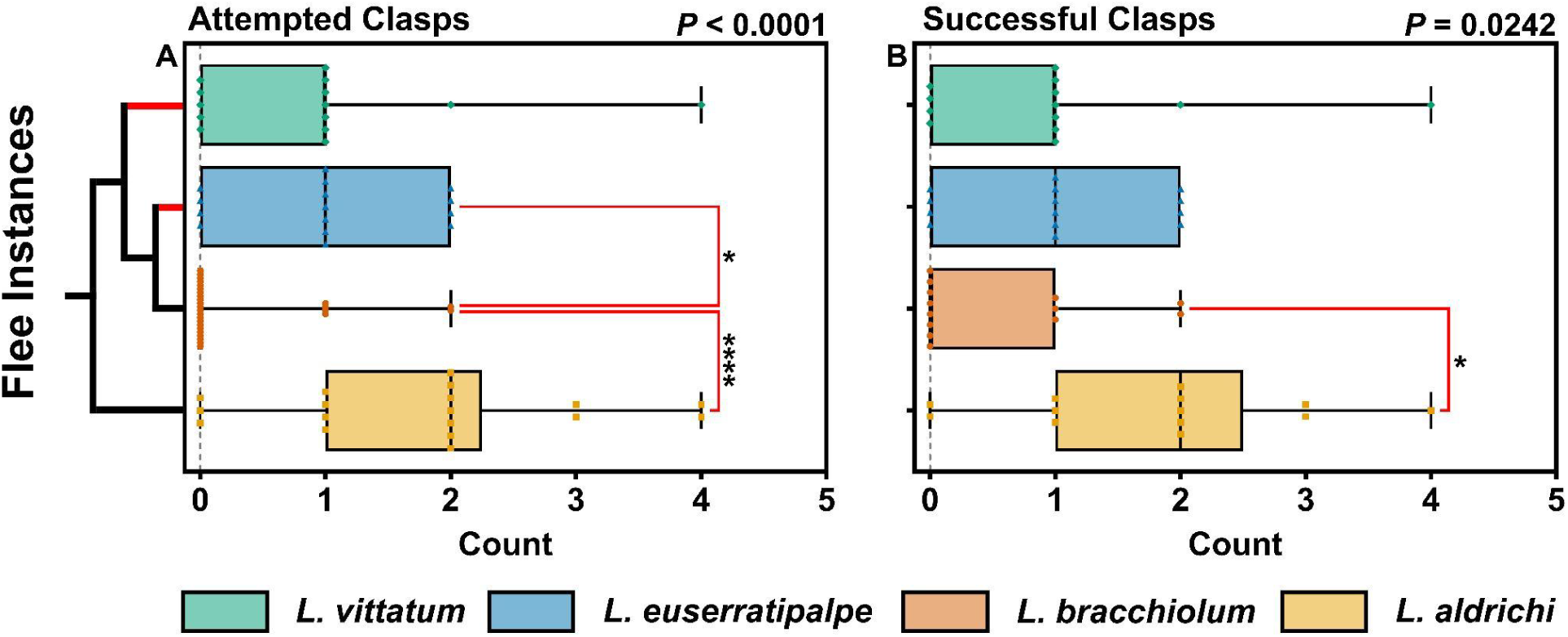
Instances of females fleeing from males during reproductive trials. Boxes represent medians plotted with IQR, symbols represent individual trial values, and whiskers correspond to the minimum and maximum trial values. The results from group comparison tests are shown in the upper right-hand corners, while significant results from post-hoc multiple comparisons tests are noted by red brackets with asterisks (* = *P* < 0.05; **** = *P* < 0.000). The simplified phylogeny indicates reproductive morphology; red tips indicate nonsacculate species and black tips represent sacculate species (phylogeny adapted from Burns et al. 2013). **A)** Attempted-clasp trials. **B)** Successful-clasp trials.

**Table A1:** Behavioural metrics recorded during this study (species codes: *Leiobunum vittatum*: Lvitt; *L. euserratipalpe*: Leus; *L. bracchiolum*: Lbr; *L. aldrichi*: Lald). Shapiro-Wilk test for normality results are shown for all metrics save average distance, which was log-normal based on a Shapiro-Wilk test for lognormality. Sample sizes for each metric follow. Note that for pursuit angle, time within one leg span, and average distance, one *L. euserratipalpe* trial could not be used due to video corruption preventing TRex scoring.

| Metric | Result | Normality | N(total) | N(Lbr) | N(Lald) | N(Lvitt) | N(Leus) |
| --- | --- | --- | --- | --- | --- | --- | --- |
| Latency | Attempts | No | 75 | 28 | 18 | 14 | 15 |
| Contact Time | Attempts | No | 75 | 28 | 18 | 14 | 15 |
|  | Successes | No | 53 | 13 | 13 | 13 | 14 |
| Pre-Attempt Contact Time | Attempts | No | 75 | 28 | 18 | 14 | 15 |
|  | Successes | No | 53 | 13 | 13 | 13 | 14 |
| Flee Count | Attempts | No | 75 | 28 | 18 | 14 | 15 |
|  | Successes | No | 53 | 13 | 13 | 13 | 14 |
| Tap Count | Attempts | No | 75 | 28 | 18 | 14 | 15 |
|  | Successes | No | 53 | 13 | 13 | 13 | 14 |
| Pursuit Angle | Attempts | No | 74 | 28 | 18 | 14 | 14 |
|  | Successes | No | 52 | 13 | 13 | 13 | 13 |
| Time within 1 Leg Span | Attempts | No | 74 | 28 | 18 | 14 | 14 |
|  | Successes | No | 52 | 13 | 13 | 13 | 13 |
| Average Distance | Attempts | Log | 74 | 28 | 18 | 14 | 14 |
|  | Successes | Log | 52 | 13 | 13 | 13 | 13 |
| Maximum Female Angle | Attempts | No | 75 | 28 | 18 | 14 | 15 |
|  | Successes | No | 53 | 13 | 13 | 13 | 14 |
| Clasping Time | Successes | No | 53 | 13 | 13 | 13 | 14 |
| Guarding Time | Successes | No | 53 | 13 | 13 | 13 | 14 |

**Table A2:** Degree of phylogenetic signal λ within the various behavioural metrics. Pagel’s lambda (Pagel 1999) was calculated using the phytools package (Revell 2012). We used a pruned version (consisting of our four study species) of a maximum clade credibility tree constructed from mitochondrial and nuclear sequences by Burns et al (2013). These models were then compared to Brownian motion predictions with a likelihood ratio test to produce our P-values.

| Metric | Trial Type | Pagel's $\lambda$ | $\log L(\lambda)$ | $LR(\lambda=0)$ | P-value |
| --- | --- | --- | --- | --- | --- |
| Latency to 1st Attempt | Attempts | 6.61e-05 | -29.451 | -6.88e-05 | 1 |
| Pursuit Angle | Attempts | 6.61e-05 | -9.263 | -2.32e-05 | 1 |
|  | Successes | 6.61e-05 | -9.831 | -3.37e-05 | 1 |
| Contact Time | Attempts | 6.61e-05 | -30.302 | -1.02e-04 | 1 |
|  | Successes | 0.999 | -28.626 | 0.187 | 0.6658 |
| Flee Count | Attempts | 6.61e-05 | -2.328 | -6.79e-05 | 1 |
|  | Successes | 6.61e-05 | -2.328 | -6.79e-05 | 1 |
| Tap Count | Attempts | 0.999 | -2.799 | 0.208 | 0.6481 |
|  | Successes | 0.999 | -2.799 | 0.208 | 0.6481 |
| Pre-Attempt Contact Time | Attempts | 6.61e-05 | -25.851 | -5.73e-05 | 1 |
|  | Successes | 6.61e-05 | -24.774 | -2.57e-05 | 1 |
| Time within 1 Leg Span | Attempts | 6.61e-05 | -29.705 | -1e-04 | 1 |
|  | Successes | 0.999 | -28.167 | 0.189 | 0.6634 |
| Average Distance | Attempts | 6.61e-05 | -6.717 | -9.68e-05 | 1 |
|  | Successes | 0.999 | -6.788 | 0.229 | 0.6325 |
| Maximum Female Body Angle | Attempts | 0.999 | -19.163 | 0.057 | 0.8111 |
|  | Successes | 6.61e-05 | -18.764 | -6.09e-05 | 1 |
| Clasping Time | Successes | 6.61e-05 | -28.053 | -1.97e-05 | 1 |
| Guarding Time | Successes | 6.61e-05 | -30.947 | -7.55e-06 | 1 |

**Table A3:**
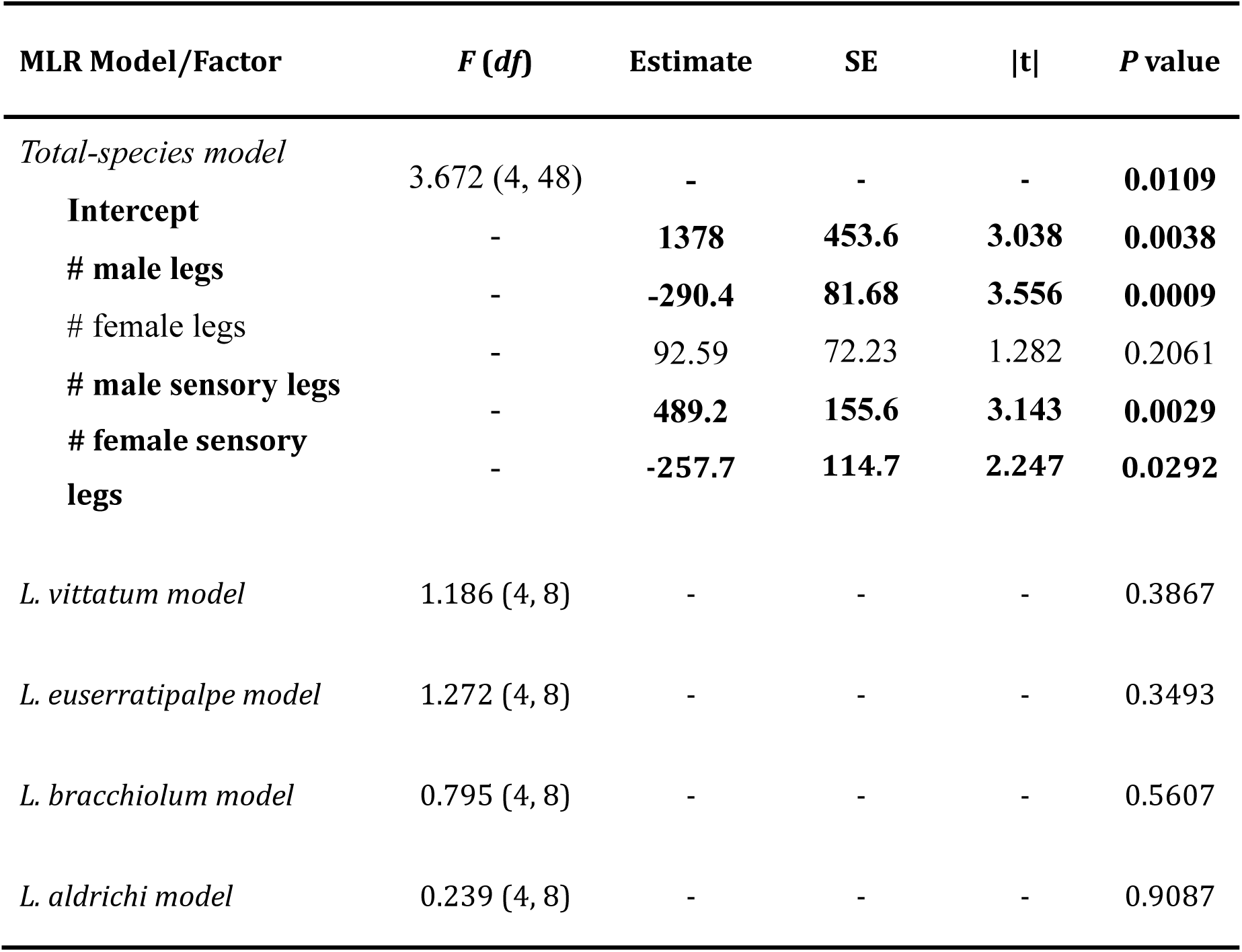
Multiple linear regression of clasping duration with leg number metrics as predictor variables for the total-species model and species-level models. Significant predictor variables are in bold; no predictor variables were dropped from modelling.

